# Epigenetic signature of human immune aging: the GESTALT study

**DOI:** 10.1101/2023.01.23.525162

**Authors:** Roshni Roy, Pei-Lun Kuo, Julián Candia, Dimitra Sarantopoulou, Ceereena Ubaida-Mohien, Dena Hernandez, Mary Kaileh, Sampath Arepalli, Amit Singh, Arsun Bektas, Jaekwan Kim, Ann Zenobia Moore, Toshiko Tanaka, Julia McKelvey, Linda Zukley, Cuong Nguyen, Tonya Wallace, Christopher Dunn, William Wood, Yulan Piao, Christopher Coletta, Supriyo De, Jyoti Misra Sen, Nan-ping Weng, Ranjan Sen, Luigi Ferrucci

## Abstract

Age-associated DNA methylation in blood cells convey information on health status. However, the mechanisms that drive these changes in circulating cells and their relationships to gene regulation are unknown. We identified age-associated DNA methylation sites in six purified blood borne immune cell types (naïve B, naïve CD4^+^and CD8^+^ T cells, granulocytes, monocytes and NK cells) collected from healthy individuals interspersed over a wide age range. Of the thousand of age-associated sites, only 350 sites were differentially methylated in the same direction in all cell types and validated in an independent longitudinal cohort. Genes close to age-associated hypomethylated sites were enriched for collagen biosynthesis and complement cascade pathways, while genes close to hypermethylated sites mapped to neuronal pathways. In-silico analyses showed that in most cell types, the age-associated hypo- and hypermethylated sites were enriched for ARNT (HIF1β) and REST transcription factor motifs respectively, which are both master regulators of hypoxia response. To conclude, despite spatial heterogeneity, there is a commonality in the putative regulatory role with respect to transcription factor motifs and histone modifications at and around these sites. These features suggest that DNA methylation changes in healthy aging may be adaptive responses to fluctuations of oxygen availability.

## INTRODUCTION

Human aging is associated with site-specific changes of DNA methylation. Summary measures of DNA methylation called “epigenetic clocks” are extensively used in aging research to estimate biological aging(1–3). Epigenetic clocks closely approximate chronological age and beyond age, predict adverse health conditions, including frailty (4), Alzheimer’s disease (5) and mortality(6, 7).

Research suggest that changes in DNA methylation with aging are regulated by specific mechanisms rather than by a stochastic drift (8). For example, a loss-of-function mutation in the H3K36 histone methyltransferase has been associated with epigenetic aging in mice (9). In humans, polymorphisms in the telomerase gene (TERT) (10) and age-dependent gain of methylation in the Polycomb repressive complex 2 have been related to accelerated aging(11). However, so far, no sound hypothesis exists that explains the association of DNA methylation with aging and pathology.

A main obstacle in understanding mechanisms driving age-associated changes of DNA methylation is that most human studies were performed in mixed blood cell types. The few studies that investigated select immune circulating cells failed to propose a unifying biological hypothesis explaining predictable changes of DNA methylation with aging(12–18).

We analyzed age-associated methylation in 6 purified blood-borne cell types sorted from peripheral mononuclear cells (PBMCs) from individuals of different ages. To minimize the confounding of age-associated pre-clinical and clinical diseases, participants were ascertained to be healthy by trained health professionals according to strict clinical criteria. We looked for CpGs differentially methylated with aging in the same direction in multiple cell types. Next, in each cell type, we conducted enrichment analyses of genes close to age-associated CpGs. Finally, we looked for chromatin accessibility markers and transcription factor binding sites close to the same age-associated CpGs. Our findings suggest that changes in methylation with aging are related to fluctuation of energetic metabolism during the life course.

## RESULTS

### Age-associated methylation in individual cell types

A principal component analysis on normalized DNA methylation (Figure 1A and Supplementary Table 1) showed that clustering by cell types was stronger than by age (PC2 showed cell types-based clustering-11.3 %) (Supplementary Fig. 1A).

**Figure 1:**
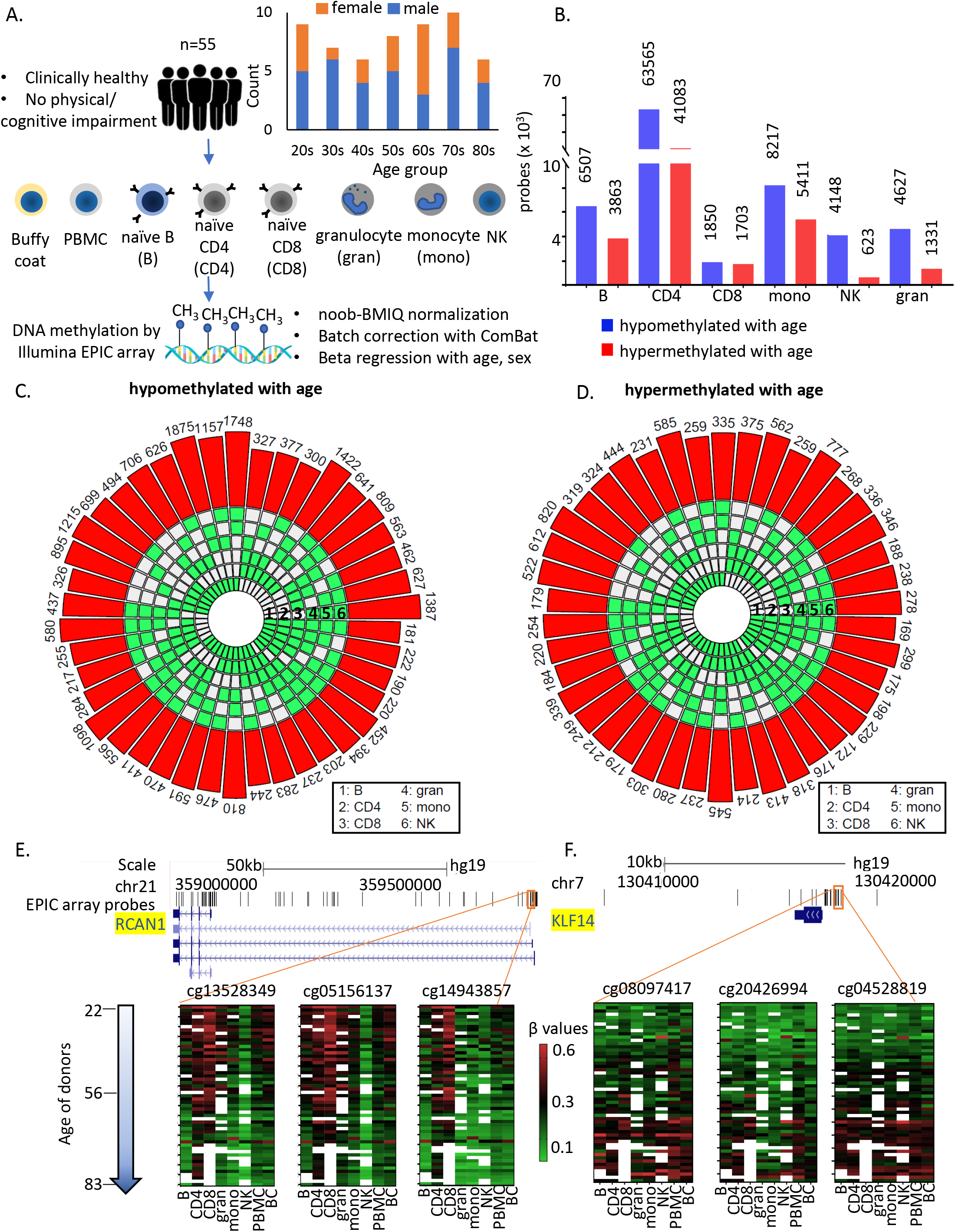
Study design and identification of age-associated methylation probes. A) Study design. B) Age-associated CpG methylation (FDR p<0.05) in 6 cell types. C-D) SuperExactTest circular plots to show the number of age-associated hypo- and hyper-methylated probes shared among different combinations of cell types (indicated by green boxes), respectively. The outermost bars show the number of probes shared among each cell type combination (regardless of other cell types). For examples, probes hypomethylated with age in B + CD4 + CD8 + gran + mono (n=222) includes probes also hypomethylated in NK cells (n=181) and probes not hypomethylated with age in NK cells (n=41). Based on the exact probability distributions of multi-set intersections, all the overlaps shown are highly statistically significant (p<10^-100^). E) Graphical representation of age-associated hypomethylation in promoter region of RCAN1 in all 6 cell types. F) Graphical representation of age-associated hypermethylation in promoter region of KLF14. The methylation status in PBMC and buffy coat are also shown. Missing methylation data is represented in white.

Age-associated CpGs were identified through sex-adjusted beta regression models (FDR corrected p-value <0.05). Number of hypo- or hypermethylated sites varied considerably between cell types (Figure 1B) with highest numbers in CD4^+^ T cells (Supplementary Fig. 1B and Supplementary Table 2). Using a different approach of comparing between young (<35 years, 25th percentile) and old (>70 years, 75th percentile) individuals, we observed >90% overlap with beta regression-derived hypomethylated sites and 70-95% overlap with hypermethylated sites in all cell types except CD8^+^ T cells (9-14% overlap) (Supplementary Fig. 1C). Having fewer old donors with CD8^+^T cells may have contributed to differences (Supplementary Table 1).

Similar to other studies, we found that a significant proportion of age-hypomethylated CpGs were in the intergenic and open-sea (>4kb from CpG island) regions while age-hypermethylated CpGs were in promoters and CpG islands (Chi sq test p<0.001) (Supplementary Figs. 1D and 1E). Additionally, age-associated differentially methylated sites in PBMC poorly recapitulate age-dependent changes that take place in specific primary immune cells (Supplementary Figs. 1E-F). These findings point to a wide heterogeneity of age-differential CpG methylation across immune blood cells and suggest that studies in PBMC poorly represents the changes that take place in specific cell types with aging.

### Shared age associated methylation across cell types

Only 181 age-associated hypomethylated sites and 169 hypermethylated sites were shared between all 6 cell types. These numbers increased to 776 (age-hypomethylated) and 404 (age-hypermethylated) sites in 5 or more cell types (Figures 1C-D). Thus, most age-related methylation changes are cell-specific. Of note, only 10 of the sites overlap with the 359 CpGs in Horvath’s pan-tissue epigenetic clock (19). While the number of shared age-hypo or hypermethylated CpGs across cells was relatively small, it was highly significantly higher than that expected based on chance alone, suggesting that common underlying epigenetic mechanisms exist across the considered cell types (Figure 1C & D). For example, CpG sites adjacent to *RCAN1* (calcineurin 1) and *KLF14* (Krueppel-Like Factor 14) show similar age-associated patterns in all cell types (Figure 1E and F).

Next, we tested whether the top 15 genes annotated to the most significant age-associated CpGs were common across multiple cell types (Figure 2A-B and Supplementary Figs. 2A-E). Only the age-hypomethylation of *CCDC102B* was common to all cell types (Supplementary Tables 3 and 4) while *ELOVL2, GPR78, LHFPL4* and *KLF14* were commonly age-hypermethylated in all cell types (Supplementary Tables 3 and 4). These findings suggest that most CpGs with age-associated methylation consistent across cell types undergo moderate (although significant) methylation changes with aging.

**Figure 2:**
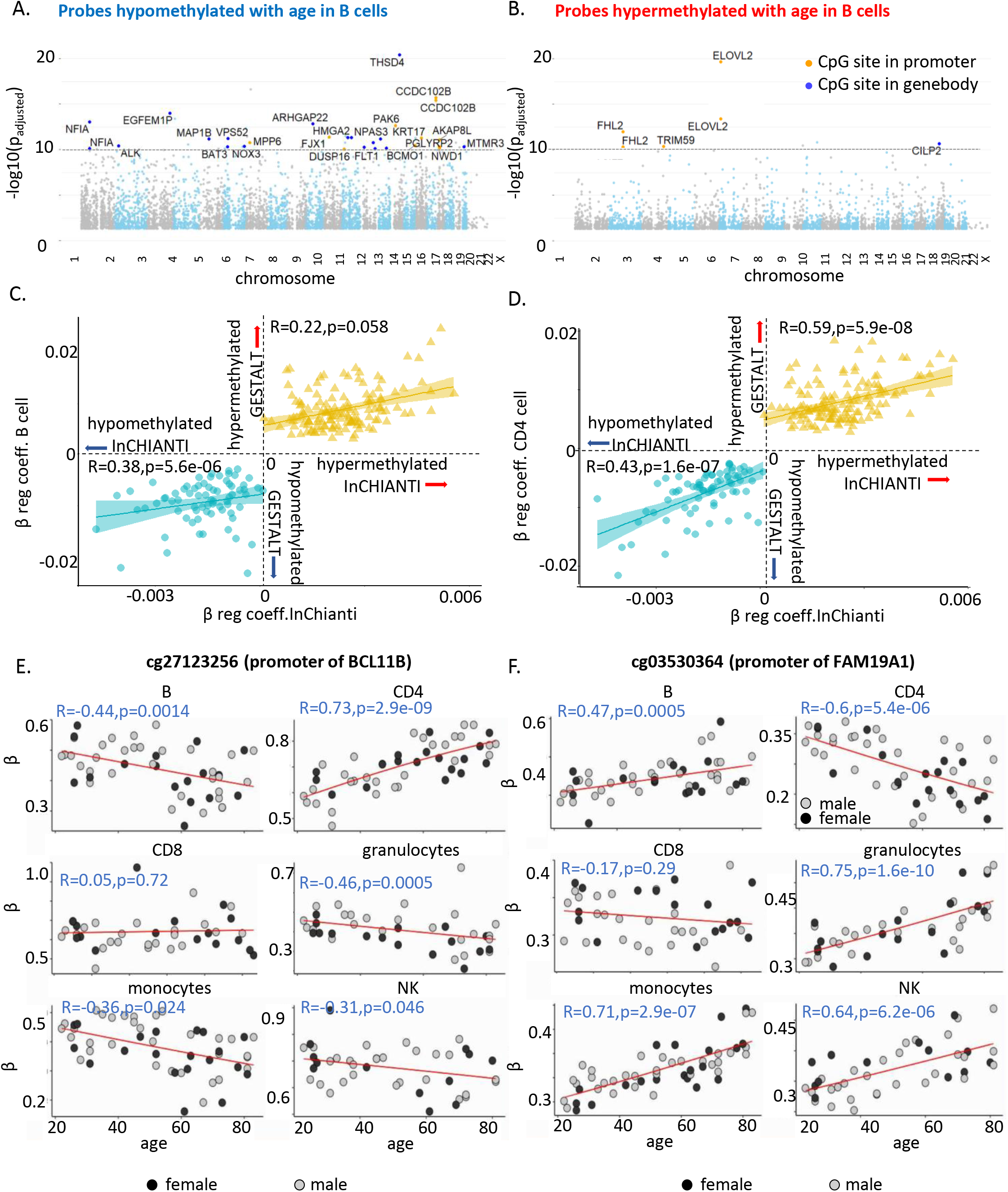
Characteristics of age-associated probes. A-B) Manhattan plot of age-associated hypo- and hypermethylated CpG sites in B cells respectively. Most significant genic probes (-log padj <10) are labelled. C) Correlation between beta-regression coefficients of age-differentially methylated CPGs in GESTALT and longitudinal InCHIANTI study. X-axis-InCHIANTI, Y-axis-B cell (Figure 2C) and CD4^+^ T cell coefficients (Figure 2D). Blue dots - age-hypomethylated CpGs, yellow triangles-age-hypermethylated CpGs. E and F) Scatter plot of age-associated CpGs showing opposite trends in different immune cells. E) cg27123256 (in BCL11B promoter) is hypomethylated with older age in B, monocytes and NK while is hypermethylated with older age in CD4^+^T cells. F) cg03530364 (in FAM19A1 promoter) is hypermethylated with older age in B, granulocytes, monocytes and NK cells while it is hypomethylated with older age in CD4^+^T cells.

### Longitudinal validation of age-associated CpG sites

We hypothesized that the age-associated CpGs identified across the six immune cells in this cross-sectional study would also show longitudinal changes of the size and direction predicted. We used DNA methylation data (Illumina 450K microarray on DNA from buffy coats) assessed at baseline and 9- and 13-year follow-up in 699 participants of the InCHIANTI study (20). Of the 181 hypo-methylated and 169 hypermethylated CpGs with age in all cell types in GESTALT, 72 and 135, respectively, were represented in the 450K microarray. The beta-coefficients for age of the 207 CpG probes (72+135) estimated from the GESTALT study and their corresponding values estimated longitudinally from the InCHIANTI study were highly and significantly correlated (hypomethylated with age CpGs: r=0.49, p=1.2e-09 and hypermethylated with age CpGs: r=0.5, p=6.9e-06 for average beta coefficients across 6 cell types, Figures 2C-D and Supplementary Figs. 3A-B). Thus, CpGs identified as differentially methylated with aging across cell types in GESTALT also change longitudinally with aging.

### Age-associated probes with opposite trends in different immune cells

Several CpGs showed significant but opposing age-trends in different cell types, especially in B, CD4^+^ T cells and monocytes (Supplementary Figs. 2F and G). For example, cg27123256 in the gene body of *BCL11B* was age-hypomethylated in non-T cells and significantly age-hypermethylated in naïve CD4^+^ T cells (Figure 2E). Our observations implicate BCL11B in aging-related changes in naïve CD4^+^ T cell function, distinct from its proposed role in effector cells (14, 21, 22). Conversely, cg03530364 in the body of FAM19A1 gene was hypermethylated in non-T cells but age-hypomethylated in CD4^+^ T cells (Figure 2F). Of note, none of these CpGs were differentially age-methylated in PBMC. Thus, opposite age-methylation trends in specific cell types may cancel each other and obscure their relevance for aging when mixed cell type sample are assessed.

### Pathway analysis of age-associated genes

Gene set enrichment analyses were performed on genes associated with at least one CpG significantly age-hypo or hypermethylated in 5 or more cell types. We identified 30 pathways (q-value<0.05) (Figure 3 and Supplementary Table 5). Probes commonly age-hypomethylated in 5 or more cell types (n=776) pointed to genes enriched in collagen biosynthesis, complement cascade and GTPase pathways (left-most column in bottom panel of Figure 3) that highlighted inflammatory and metabolic pathway in aging. Genes associated with shared age-hypermethylated probes (n=404) were enriched for neural pathways previously implicated to brain aging along with G-Protein Coupled Receptors pathways (23) (left-most column in top panel of Figure 3). Key pathways are highlighted, with associated genes displayed in boxes on the right-hand side.

**Figure 3:**
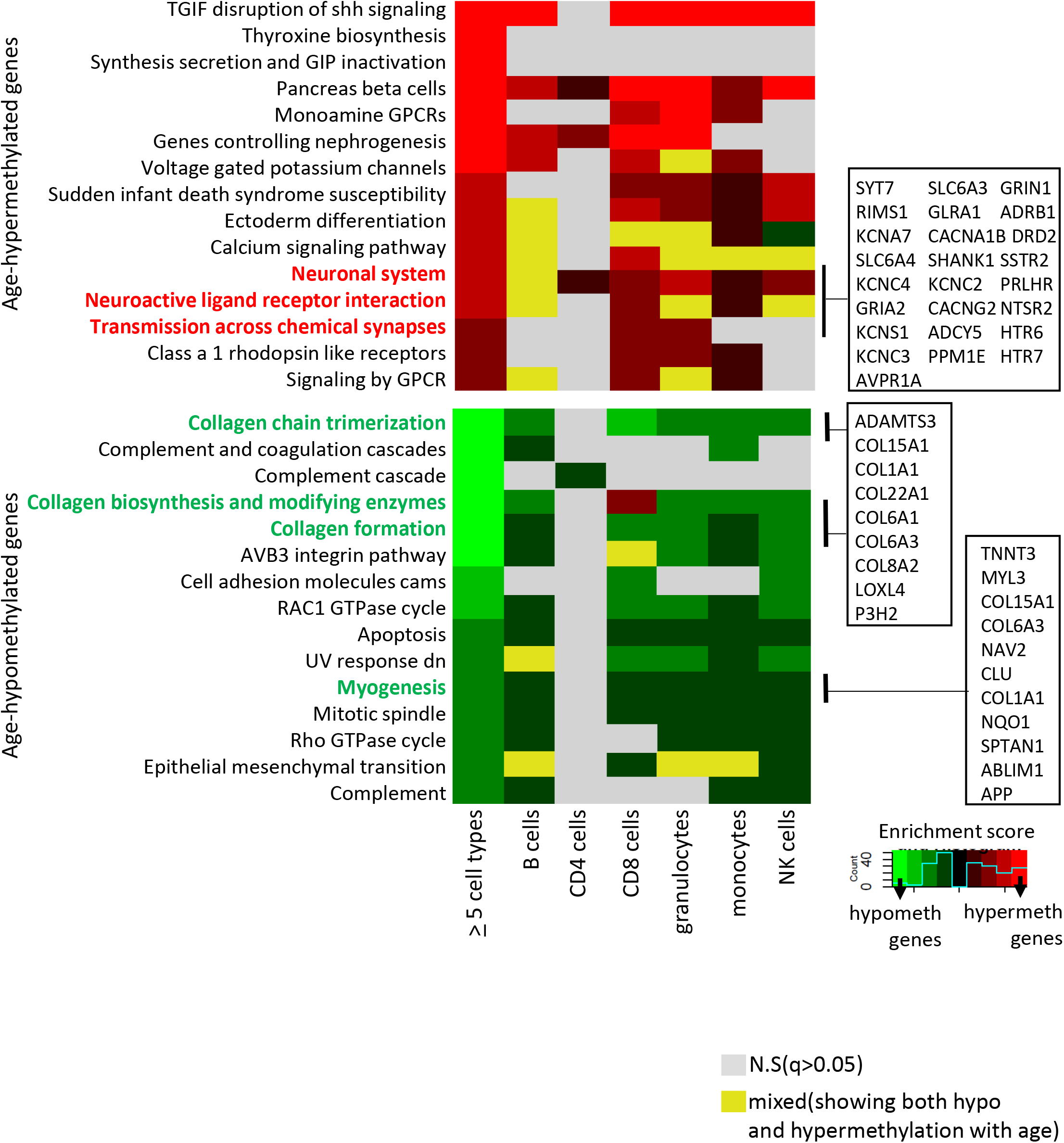
Pathway analysis of methylated probes. Enrichment analysis of genes annotated to age-associated hypo- and hyper-methylated CpGs in ≥5 cell types (left-most column) and in individual cell types. Red/green shades indicate enrichment scores in hyper- (red) and hypo- (green) methylated genes. Yellow indicates ambiguous pathways associated with both hypo- and hyper-methylated genes in individual cell types. Not significant pathways are shown in grey. Full results in Supplementary Table 5.

### Functional annotation of age-associated probes

To further interrogate the relationships between DNA methylation and other epigenetic states, we mapped the methylation age-associated sites to cell-specific chromHMM-derived chromatin profiles(24). As controls, we annotated all sites in the EPIC array to the 18-state chromHMM model of respective primary cell type. Granulocytes were excluded from this analysis because reference data were not available.

Age-associated hypomethylated CpGs were significantly enriched for weak/active enhancers (yellow bar, Figure 4A) whereas, confirming previous reports, age-hypermethylated CpGs, were enriched in bivalent/polycomb regions compared to control set (brown and dark grey bars respectively in Figure 4A). Results for cell types-specific analyses are shown in Figure 4B.

**Figure 4:**
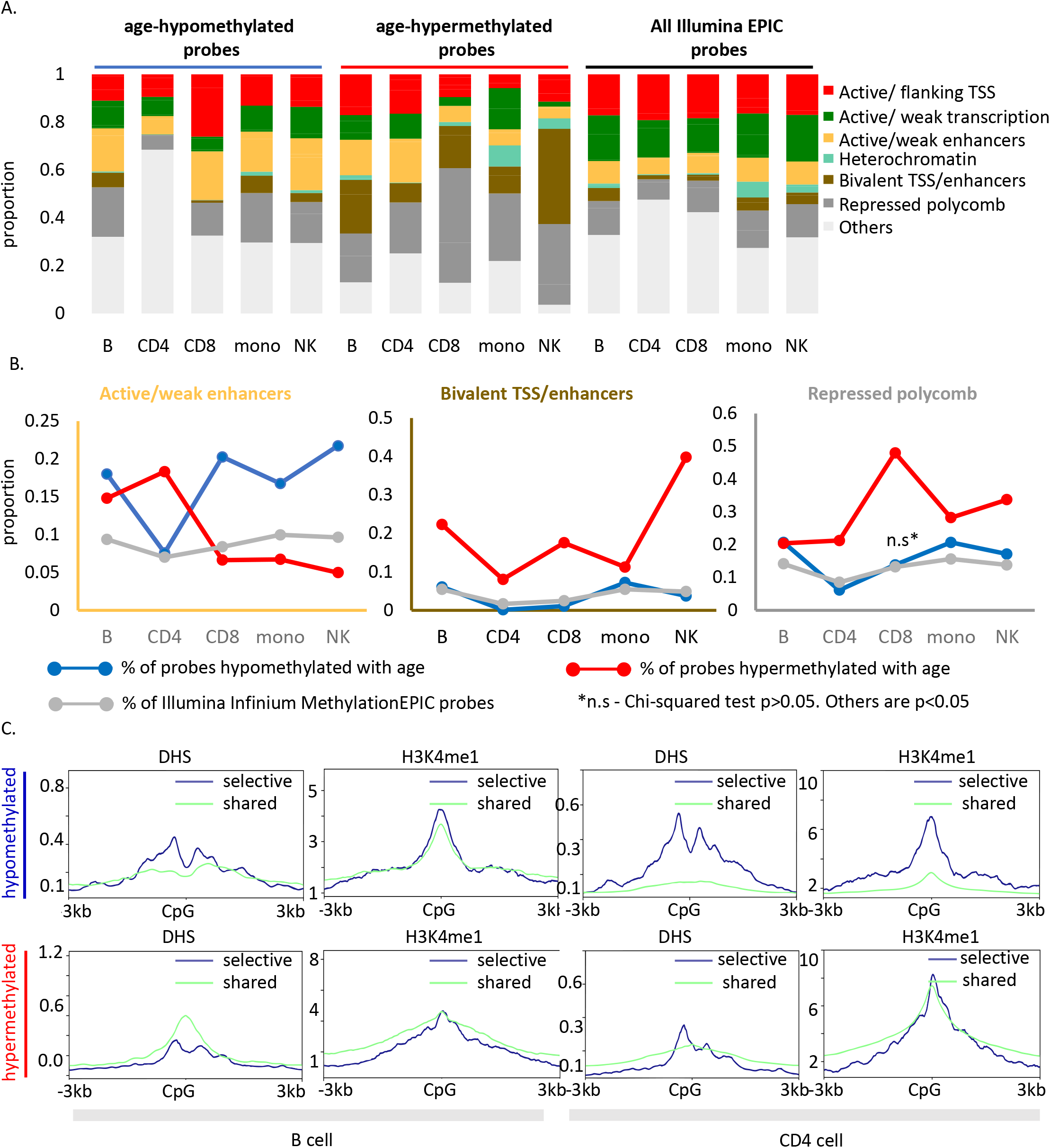
Functional annotation of age-associated probes along with their grouping based on sharedness. A) ChromHMM annotation of age-associated CpGs. B) Proportion of CpGs mapping to weak/active enhancers (left, orange box), bivalent enhancers/TSS (inset, brown box) and polycomb repressor regions (right, grey box) in age-associated hypo- (blue line), hypermethylated (red line) CpGs as compared to all MethylationEPIC CpGs (grey line). C) DeepTools plots showing the distribution of accessible chromatin (DNase hypersensitive sites) and H3K4me1 histone mark in and around ± 3kb region of age-differentially methylated CpGs. The age-associated sites were divided into shared (blue) (common between 5 or more immune cells) and selective sites (green). The top row shows the pattern for age-associated hypomethylated CpGs while the bottom row is for the age-associated hypermethylated CpGs in B and CD4^+^ T cells.

We further mapped the profile of four epigenetic markers from the ENCODE project in and around (+ 3kb) age-associated methylation sites. For B and CD4^+^ T cells, we observed a V-shaped peak-valley-peak pattern of DNase hypersensitivity at sites of age-associated hypomethylation, which is characteristic of promoter sites (Figure 4C) (25). Both age-associated hypo- and hypermethylated sites showed evident H3K4me1 peaks, a marker commonly associated with active and primed enhancers (Figure 4C)(26). No specific trend was observed for H3K4me3 and H3K27ac (data not shown). These patterns were highly consistent across cell types (Supplementary Fig. 4) and strongly suggest functional connections between methylation and chromatin status.

### Pattern of transcription factor binding motifs around age-associated CpGs

Specific transcription factors (TF) may induce or been induced by DNA methylation (27, 28). Through our *de-novo* HOMER analysis, we observed that the binding motif for aryl hydrocarbon receptor nuclear translocator (*ARNT*, also named HIF1β) was associated with age-hypomethylated CpGs across most cell types (Figure 5A). The only exception was naïve CD8^+^ T cells where the top enriched motif was B-cell lymphoma gene 6(*BCL6*). *BCL6* code for a zinc finger transcription factor that plays a critical role in the generation of memory and effector cells in acute infection (29). Another motif associated with age-hypomethylated CpGs across most cell types was chromatin architectural protein CTCF and its closely related gene BORIS. Methylation changes at CTCF sites reflected large scale genome reorganization in immune cells in older individuals (30, 31).

**Figure 5:**
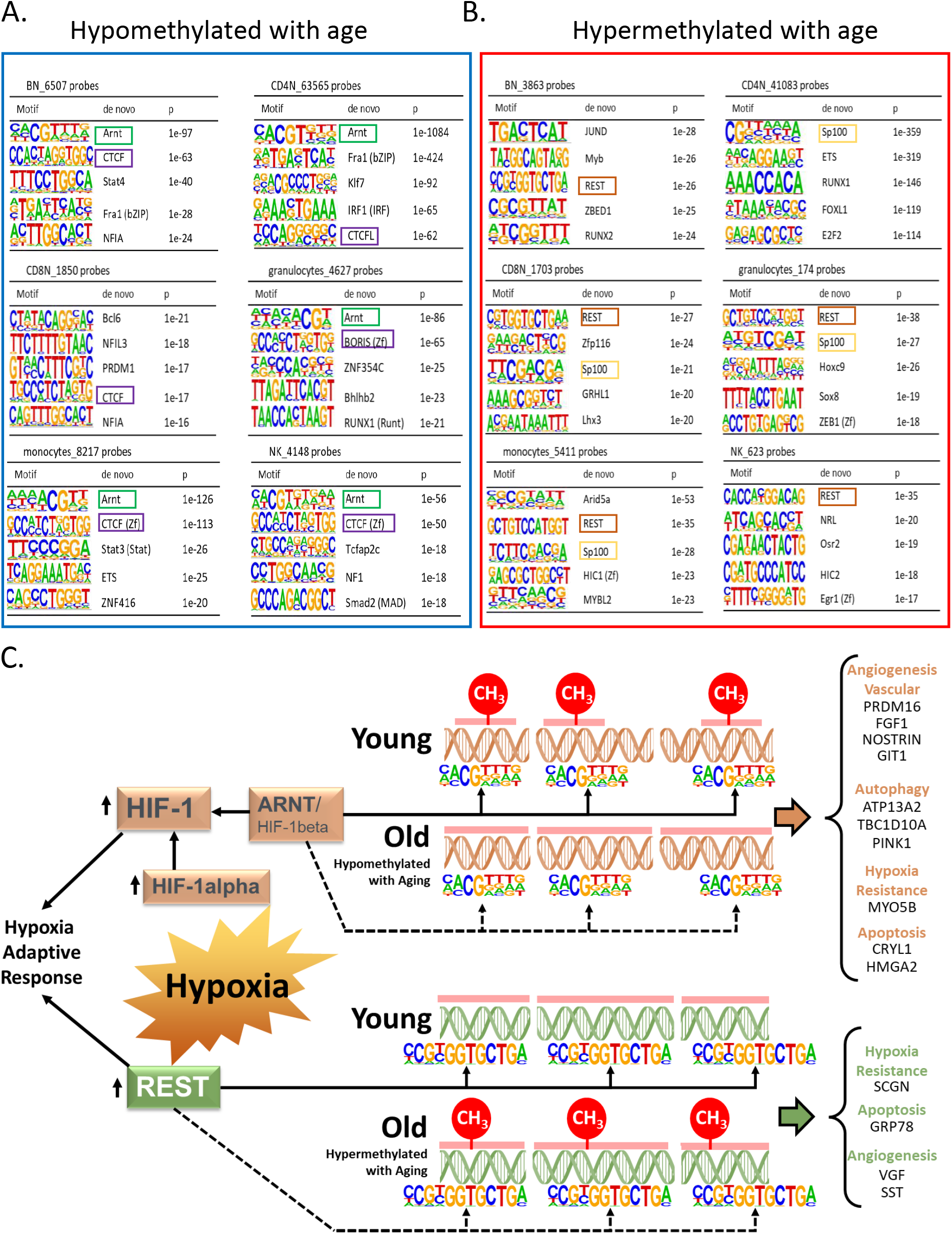
Association of transcription factor binding motifs with age-differentially methylated CpGs. A) Top 5 TF motifs at and around (+ 200bp) of CpG sites that are hypomethylated with age. Recurring motifs like ARNT and CTCF/BORIS are highlighted. B) Top 5 TF motifs at and around (+ 200bp) CpG sites that are hypermethylated with age. Recurring motifs like REST and Sp100 are highlighted. C) Hypoxia-centric model of age-associated sites with ARNT and REST motifs. CpG sites hypomethylated with aging across 6 different cell types are significantly more likely to host binding motifs for ARNT, the core hub for the hypoxia response. On the contrary, CpG sites hypermethylated with aging are significantly more likely to host binding motifs for REST, a hypoxia response transcriptional repressor. On the right are selected age-associated genes that carry the motifs for ARNT or REST transcription factors.

Repressor Element 1-Silencing Transcription Factor (*REST*) was the TF motifs most frequently associated with age-hypermethylated CpGs in 5 of 6 cell types (Figure 5B). Age-hypermethylated sites in PBMCs have been previously shown to be enriched for *REST,* which is known to repress stress response genes and is lost in cognitive impairment and Alzheimer’s disease pathology (32, 33). The top enriched TF motif associated with age-hypermethylated sites in monocytes was Arid5A (p<10^-27^) that binds to selective inflammation-related genes, such as *IL6* and *STAT3* and stabilize their expression (34, 35).

The recurring enrichment of *ARNT* and *REST* with age-associated CpGs observed across multiple cell types, despite relatively few shared genomic region locations, suggests this relationship is functional. We found that only 17 and 44 age-associated hypo- and hypermethylated probes, respectively, shared *ARNT* or *REST* motifs across all cells (Supplementary Figs. 5A-B), suggesting these overlaps are not random and have a specific function (Supplementary Figs. 5A and B).

Remarkably, *ARNT* was significantly overexpressed in older age in three of the six cell types and REST showed a significant decrease of expression with age in most cell types (Supplementary Table 6). These findings suggest that age-associated changes in expression levels of *REST* and *ARNT* can affect the epigenetic status of their target genes.

### Age-related differential methylation and oxygen sensing

*ARNT, REST* and *BCL6,* three transcription factors most associated with differentially methylated regions, are implicated in hypoxia response (Figure 5C). *ARNT* is the beta subunit of Hypoxia Factor 1 (HIF-1), which is stabilized during hypoxia and shuttled to the nucleus where it binds to DNA hypoxia response elements (HRE) and triggers a complex response that include upregulation of angiogenesis and erythropoiesis and reprogramming of energetic metabolism from oxidative phosphorylation to anaerobic glycolysis (36). Hypoxia also upregulates the transcription of *REST* which is the master regulator of the transcriptional repression arm of the response to hypoxia. Released REST is shuttled to the nucleus where it binds to DNA and regulates approximately 20% of the hypoxia-repressed genes, including genes involved in proliferation, translation, and cell cycle progression. We identified 35 genes that were hypomethylated with aging and had close by an ARNT motif in all six cell types (Data not shown). Ten of these genes (right side of Figure 5C, genes under orange headings) have been linked to hypoxia response (37–46). Similarly, we found 20 genes with probes hypermethylated with age and with REST motif in the vicinity in all six cell types (data not shown). Four of these (right side of Figure 5C, genes under green heading) are known to be downregulated in hypoxia (47–50). These results strongly suggest a link between age-associated DNA methylation and oxygen sensing through putative regulation by transcription factors like *ARNT* and *REST* in the various immune cells.

## DISCUSSION

Novel and important conclusions arise from our observations. First, only few CpG sites are hypomethylated and hypermethylated with aging across all circulating cells while the majority of significant age-associated methylation changes are cell-selective. Indeed, several CpGs show differential age-methylation in opposite directions in different cell types and are unchanged in PBMC, suggesting that they may be missed when studying mixed cell samples. Noteworthy, age-related methylation differences in this crosssectional study were strongly and significantly correlated with longitudinal age-associated methylation changes in an independent population.

Age-associated hypomethylated sites were significantly enriched for active enhancers whereas age-hypermethylated sites were enriched for bivalent/polycomb regions, confirming previous findings in whole blood(32). Age-differential methylation coincided with specific chromatin status and histone markers patterns, suggesting that their position in proximity of promoter and active enhancer regions is connected with chromatic accessibility and potentially modulation of gene expression.

Third, distinct TF binding motifs co-localize with CpGs differentially methylated with aging despite wide variation in the distribution of such sites across cell types, suggesting a specific regulatory function. Noteworthy, the top age-associated TF identified, *ARNT* and *REST* act in coordination in hypoxia response (51). *BCL6,* another top TF binding motif associated with age-differentially methylated CpG has also been shown to protects cardiomyocyte from damage during hypoxia (52). These finding supports the hypothesis that systematic methylation changes with aging may be induced by fluctuations in oxygen availability and energy metabolism. Interestingly, the mRNA encoding *ARNT* significantly increases with age in all cell types except monocytes, while mRNA coding for *REST*declines with aging in 4 cell types and shows no significant change in naïve CD8+ T cells and NK cells. mRNAs coding for *CTCF* showed strong age-association across numerous cell types (Supplementary Table 6). The hypothesis that oxygen sensing regulates directly or indirectly DNA methylations is consistent with studies showing that in replicating fibroblasts, biological age estimated by DNA methylation slows down under hypoxia compared to normoxia(53). Further, many genes close by to “shared” age-differentially methylated CpG identified in our analyses play important roles in hypoxia response (Figure 5C).

The specific mechanisms connecting age-related changes in DNA methylation in genes which also contain binding motifs the master hypoxia-response mediators remain unknown. Shahrzad al. reported an inverse correlation between the severity of hypoxia and the degree of DNA methylation(54). There is evidence that hypoxia-induced hypermethylation may be due to reduced TETs activity (55). Our findings add to this literature by suggesting that a direct interaction between hypoxia-related transcription factors and DNA methylation at specific DNA sites occur with aging, perhaps as an adaptive response triggered by fluctuations in oxygen levels that occur in many age-related conditions. This hypothesis is consistent with oxygen availability been the most important environmental factor that requires physiological adaptation during pregnancy and development and extends this concept in a life course perspective.

A limitation of this study is that we have focused on circulating cells and, therefore, our findings may not apply to age-methylation in other tissues. In addition, our findings were not replicated in an independent cross-sectional study population. In spite of these limitations, this study has unique features: a cohort of exceptionally healthy donors and percent methylation was assessed in specific cell types obtained by cytapheresis and sorted by using state-of-the art methods.

## CONCLUSION

Age-associated DNA methylation profiles of the six purified primary immune cell populations in the blood show more cell-specificity than sharedness. However, we observe common regulatory features with respect to transcription factor binding motifs and histone modifications. Based on the consistent association of these methylated sites with ARNT and REST, which are master hypoxia regulators, we hypothesize that oxygen sensing and hypoxia drive mechanisms for changes in methylation. This hypothesis should be further explored in animal models with manipulation of oxygen levels and serial measures of DNA methylation in circulating immune cells.

## MATERIALS AND METHODS

### Cohort details

Buffy coat, peripheral blood cells (PBMC) and granulocytes were collected from Genetic and Epigenetic Signatures of Translational Aging Laboratory Testing study (GESTALT) study participants (N=55; 34 men and 21 women; age 22-83 years) who were free of diseases (except controlled hypertension or history of cancer silent for > 10 years), not on medications (except one antihypertensive drug), had no physical or cognitive impairments, non-smokers, weighed > 110 lbs, had BMI < 30 kg/m^2^ (56) (57). GESTALT was approved by the institutional review board of the National Institutes of Health and participants explicitly consented to participate.

### Isolation of PBMC and immune cell populations

PBMCs were isolated from cytapheresis packs by density gradient centrifugation using Ficoll-Paque Plus. Total B, CD4^+^ and CD8^+^ T cells were enriched by negative selection using EasySep Negative Human kits specific for each cell type; monocytes were negatively enriched using “EasySep Human Monocyte Enrichment Kit w/o CD16 depletion”. Natural killer cells were negatively enriched by depleting PBMCs with antibodies against CD3, CD4, CD14, CD19 and Glycophorin-A in HBSS buffer. Enriched cell populations were FACS sorted by flow cytometry as per Human Immunophenotyping Consortium (HIPC) phenotyping panels (58). Gating strategies and post-sort purity were analyzed by FlowJo software (LLC, Ashland, OR)(56). Granulocytes were positively selected from whole blood using EasySep™ Human Whole Blood CD66b Positive Selection Kit. Purified cells and PBMC were washed with PBS, snap frozen and stored at −80° C. All sorted cells were >95% pure by flow cytometry(56).

### Assessment of DNA methylation

DNA was isolated from 1–2 million cells using DNAQuik DNA Extraction protocol and the Qiagen DNeasy Kit. 300 ng of DNA was treated with sodium bisulfite using Zymo EZ-96 DNA Methylation Kit. The methylation of ~850,000 CpG sites was determined using Illumina Human MethylationEPIC BeadChip, and data preanalyzed by GenomeStudio 2011.1.

### Data processing and functional annotation of CpG sites

Analyses was performed by the R minfi package (59, 60). Probes with low detection p-values (cutoff 0.01) were filtered out(61). Data was normalized using noob and BMIQ(62), batch corrected by ComBat function (sva package), and β values were used for differential methylation analyses. Following the MethylationEPIC probe annotation (IlluminaHumanMethylationEPICanno-.ilm10b2.hg19) to the UCSC RefSeq genes (hg19), we grouped the locations into 3 categories - 1) promoter group-TSS1500 (from 201-1500 bp upstream of TSS), TSS 200 (≤200 bp upstream of TSS), 5’UTR, first exon; 2) genebody-exons (all exons except exon1), exon intron boundary, intron and 3’UTR and 3) intergenic probes. The first gene in the annotation package was considered. Probes were divided into 3 groups-within CpG islands (CGI), within CpG shore (0-2kb from CGI), CpG shelf (2-4kb from CGI) and open sea (>4kb from CGI).

### Definition of age-associated probes

Age and sex adjusted CpG-specific beta regressions were performed on normalized β values using the R *betareg* function. P-values were adjusted for multiple testing (Benjamini-Hochberg (BH) adjusted p< 0.05). Probes with FDR p<0.05 for age were considered age-differentially methylated CpGs and considered hypo-(Estimate_age_ < 0) or hypermethylated (Estimate_age_ >0). The overlap of probes across multiple combinations of the six cell types was assessed using R package SuperExactTest (v.1.1.0) (63).

### Gene Set Enrichment Analysis (GSEA)

Based on the EPICarray annotation, genes were classified as differentially hypo- or hyper-methylated with age. Genes with both age hypo- and hyper-methylated CpGs were removed from the analysis. Enrichment analysis was performed by the tmodHGtest method in the tmod v.0.46.2 R package, comparing a foreground list of genes found in ≥5 cell types against reference gene set collections “Hallmarks” and “Canonical Pathways” (which includes Reactome, KEGG, WikiPathways, PID, and Biocarta gene sets) from the Molecular Signature Database MSigDB (v.7.4)(64).

### Visualization of histone peaks and DHS peaks

Primary cell DHS and chromatin ChIP-Seq bigwig files were downloaded from ENCODE (https://www.encodeproject.org/) (56). DeepTools was used to visualize DHS and histone peaks in +3kb region surrounding age-associated shared and non-shared methylated sites. For plotting purposes, the order of methylated probes was determined based on descending score of DHS peaks and followed for all histone marks (H3K4me1, H3K4me3 and H3K27ac).

### Annotation of age-associated methylated probes using chromHMM

The 18-state chromHMM models (based on 6 chromatin marks H3K4me3, H3K4me1, H3K36me3, H3K27me3, H3K9me3 and H3K27ac) for various immune cells (E032-primary B cell, E038-primary naïve CD4^+^ T cells, E047-primary naïve CD8^+^ T cells, E029-monocyte, E046-NK cell) were downloaded from Roadmap epigenomics project (https://egg2.wustl.edu/roadmap/web_portal/chr_state_learning.html). Bedops tool was used to map the age-associated methylated sites to the respective chromHMM profiles. All Infinium MethylationEPIC array probes were also partitioned using each of the immune cell chromHMM profiles as controls.

### Prediction of de-novo transcription factor binding motifs by HOMER

+200bp around each age-associated methylated site was provided as input for analysis in HOMER using de novo setting(65).

### InCHIANTI longitudinal study cohort

InCHIANTI (Invecchiare in Chianti) is a population-based cohort of individuals ≥ 20 years old from the Chianti region of Tuscany, Italy (PMID: 11129752). The Italian National Institute of Research and Care on Aging Institutional Review Board approved the study protocol and all participants explicitly consented to participate. DNA methylation from 699 participants (1841 observations) were used for the analysis. CpG methylation of 485,577 CpGs was determined by the Illumina Infinium HumanMethylation450 BeadChip (Illumina Inc., San Diego, CA) and data processed by the R package “sesame”. Mean rates of change were estimated from 2-3 longitudinal timepoints.

### RNA-Seq sample extraction, processing and data analysis

Total RNA was extracted from 2 x10^6^ cells, depleted from ribosomal RNA and 50ng was used for cDNA synthesis and library preparation. Libraries were sequenced for 138 cycles on Illumina HiSeq 2500. After adapter removal and end trimming of raw FASTQ files, transcript abundances were quantified with reference to hg19 transcriptome using kallisto 0.44 (with options --single -l 250 -s 25). Transcripts were aggregated to genes with tximport and filtered out if less than 10TPM were detected in more than 33% of the samples. Linear regression models (~ phase + age*sex) were used on TPM normalized expression values to study expression changes of selected transcription factors with age. female.

### Data sharing statement

Microarray data are available at GEO under accession number GSE184269.

## Acknowledgements

This work was supported entirely by the Intramural Research Program of the National Institute on Aging. We are grateful to the GESTALT participants and the GESTALT Study Team at Harbor Hospital and NIA.

## Author Contributions

Conception or design of the work- N.P.W., R.S., L.F

Acquisition, analysis, or interpretation of data- R.R., P-L.K., J.C, D.S., C.U-M., D.H., M.K., S.A., A.S., A.B., J.K., A.Z.M., T.T., J.M., L.Z., C.N., T.W., C.D., R.W., W.W., Y.P., K.G.B., C.C., S.D., J.M.S.

Drafting or substantively revising the manuscript– R.R., R.S., L.F.

## SUPPLEMENTARY FIGURE LEGENDS

**Supplementary Figure 1:**
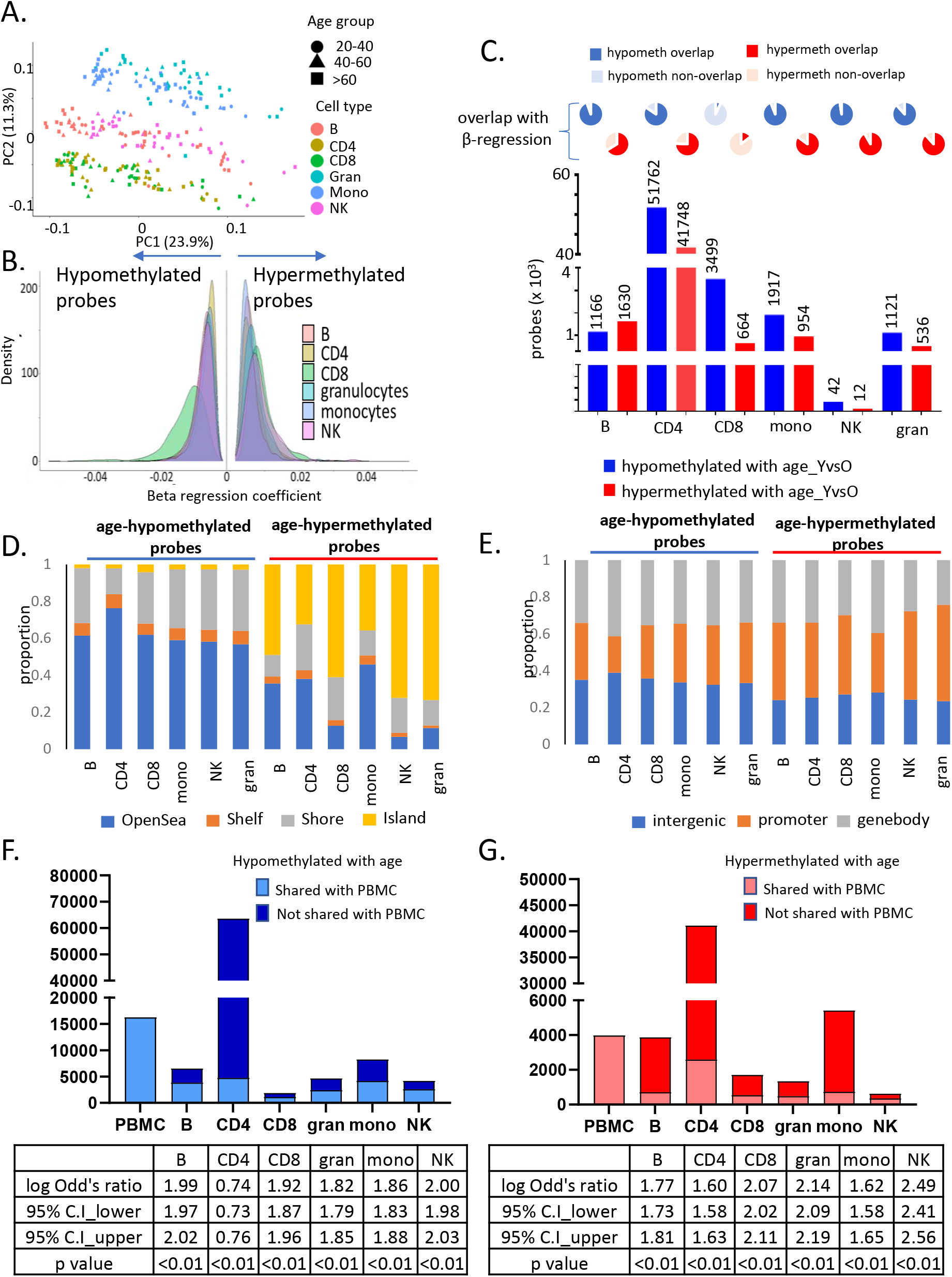
Characteristics of entire dataset and age-associated methylation data in 6 primary immune cells. A) PCA plot of normalized methylation data of six immune cells in the 55 healthy donors. The cell types are indicated in different colors, while the three broad age groups (20-40, 40-60 and 60-90 years) are indicated in different shapes (PC1-principal component 1, PC2-principal component 2). B) Distribution of beta-regression coefficient of the age-associated hypo- and hypermethylated probes in all six immune cell types estimated this cross-sectional study. Distribution of coefficient values categorized into groups is shown in Supplementary Table 2. C). Age-associated probes from independent sample t-test analysis of young (<=35years, 25th percentile) vs old (>=70years, 75th percentile) donors for each cell type. The pie charts on top show the extent of overlap with results from the beta-regression analysis. D) Distribution of age-associated probes from beta regression into groups based on distance from CpG islands (CGI) (Island-within CGI, shore-within 2kb of CGI, shelf-2-4kb of CGI, open sea->4kb from CGI). E) Distribution of age-associated hypo- and hypermethylated probes with respect to location (promoter-1500TSS to 1st exon, genebody-within exons, introns and 3’UTR). F and G) Overlap of age-associated hypo- and hypermethylated probes in the 6 immune cell types with those identified in PBMCs. The first bar indicates the number of age-associated probes identified in PBMC. The following bars show the counts in the other immune cells, the lighter portion of the bars show the number of probes that are shared with PBMCs, and the darker portion indicates non-PBMC cell-specific probes. The tables below show the log of odd’s ratio and its 95% confidence interval to represent of the significance of the overlap between age-associated probes in each cell type with PBMC.

**Supplementary Figure 2:**
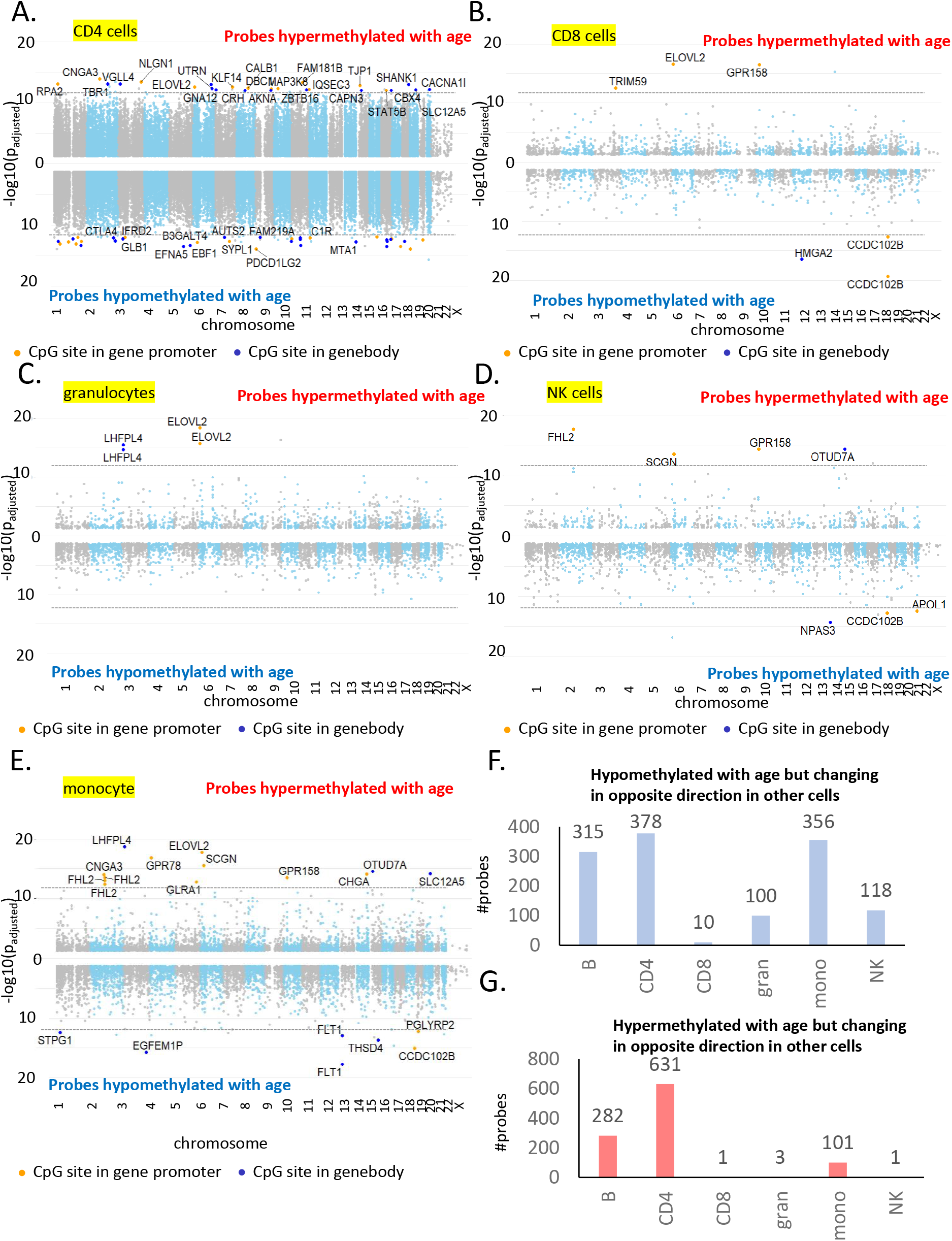
Most significant age-associated CpGs in non-B immune cells along with CpGs showing opposite age-associated trends. A) Manhattan plot of age-associated CpGs in CD4^+^ T cells. The X-axis shows the distribution of significant CpGs (FDR p<0.05), and the Y-axis shows the associated negative log of the adjusted p value from beta regression. Positive axis comprises of probes hypermethylated with age while the negative axis shows age-associated hypomethylated probes. The top hits in each group with the most significant p-values are labelled where the orange dot present CpG probes in the gene promoter. B) Manhattan plot of age-associated probes in CD8^+^T cells. C) Manhattan plot of age-associated probes in granulocytes. D) Manhattan plot of age-associated probes in NK cells. E) Manhattan plot of age-associated probes in monocytes. F) Count of probes hypomethylated with age in cell type of interest but showing hypermethylation in one or more other cell types. For example, 315 probes are age-hypomethylated in B cells but are significantly hypermethylated with age in one or more other immune cell types. Maximum number of such probes are observed in CD4^+^ T cells followed by B cells and monocytes. G) Count of probes hypermethylated with age in cell type of interest but showing hypomethylation in one or more other cell types. For example, 282 probes are hypermethylated in B cells but are significantly hypomethylated with age in one or more other immune cell types. Maximum number of such probes are observed in CD4^+^ T cells followed by B cells and monocytes.

**Supplementary Figure 3:**
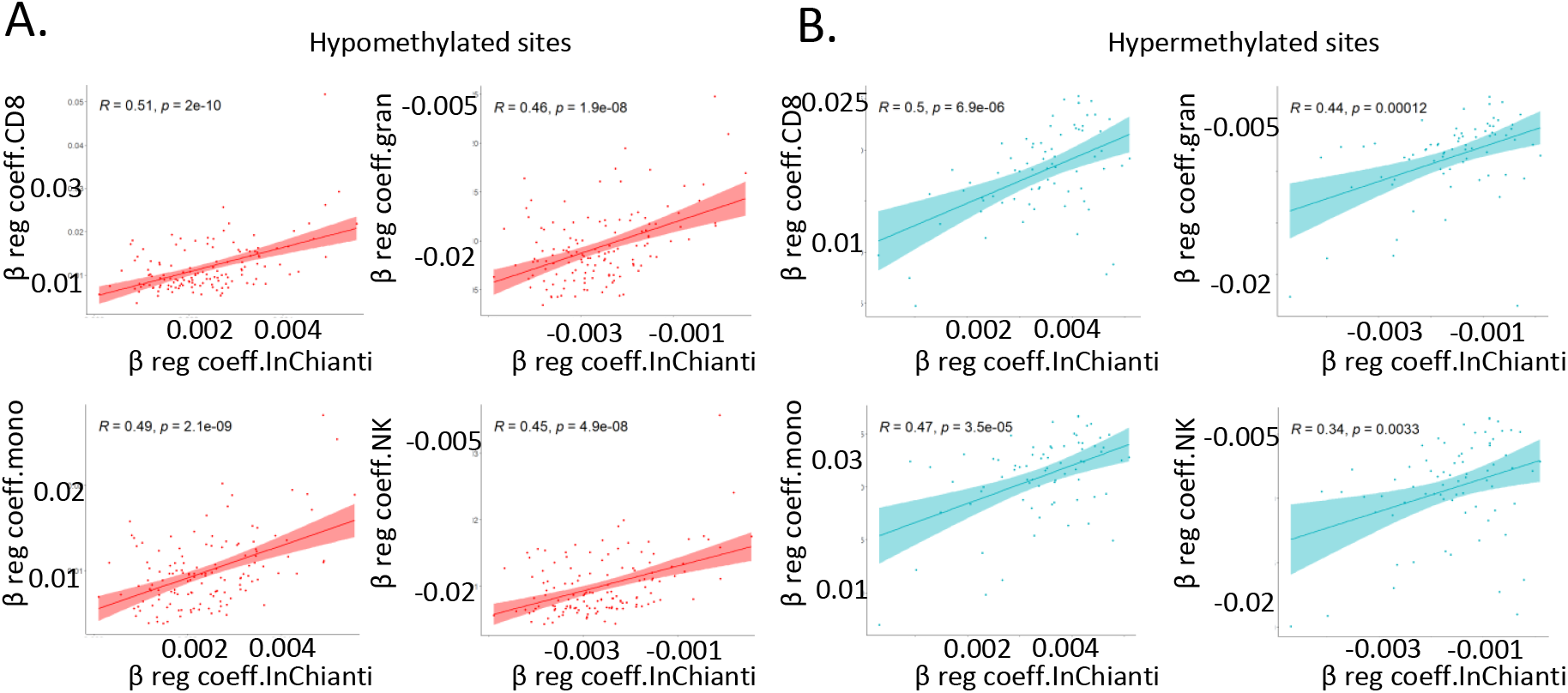
Comparison individual immune cells with InCHIANTI longitudinal study. A-B) Correlation between beta-regression coefficients of age- associated methylation probes in 5 or more cell types in study and beta-regression coefficients estimated from longitudinal data in the InCHIANTI study. On the X-axis is the data from InCHIANTI longitudinal study cohort while on the Y-axis is cell-specific coefficient values for CD8^+^ T cells (top left), granulocytes (top right), monocytes (bottom left) and NK cells (bottom right). pink dots are the coefficients of the age-hypomethylated probes (Supplementary Figure 3A) while the blue dots are for age-hypermethylated probes (Supplementary Figure 3B).

**Supplementary Figure 4:**
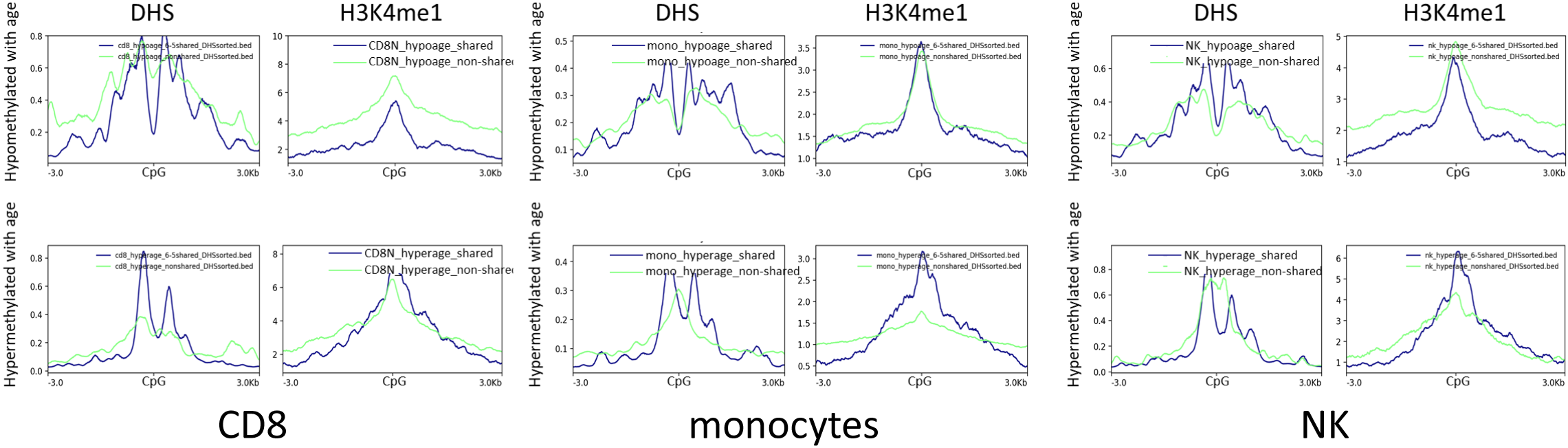
Functional annotation of age-associated probes with respect to DHS and histone marks from ENCODE. The functional significance of age-associated probes with respect to other epigenetic marks was examined using the DNase hypersensitivity sites and H3K4me1 peaks on primary immune cells from ENCODE. Granulocyte data was not available and hence could not be examined. Age-associated hypo- or hypermethylated probes were grouped into shared (blue) and selective (green) based on whether they were common across 5 or more of the 6 cell types. A region of 3kb of either side of the CpG probes of interest was examined.

**Supplementary Figure 5:**
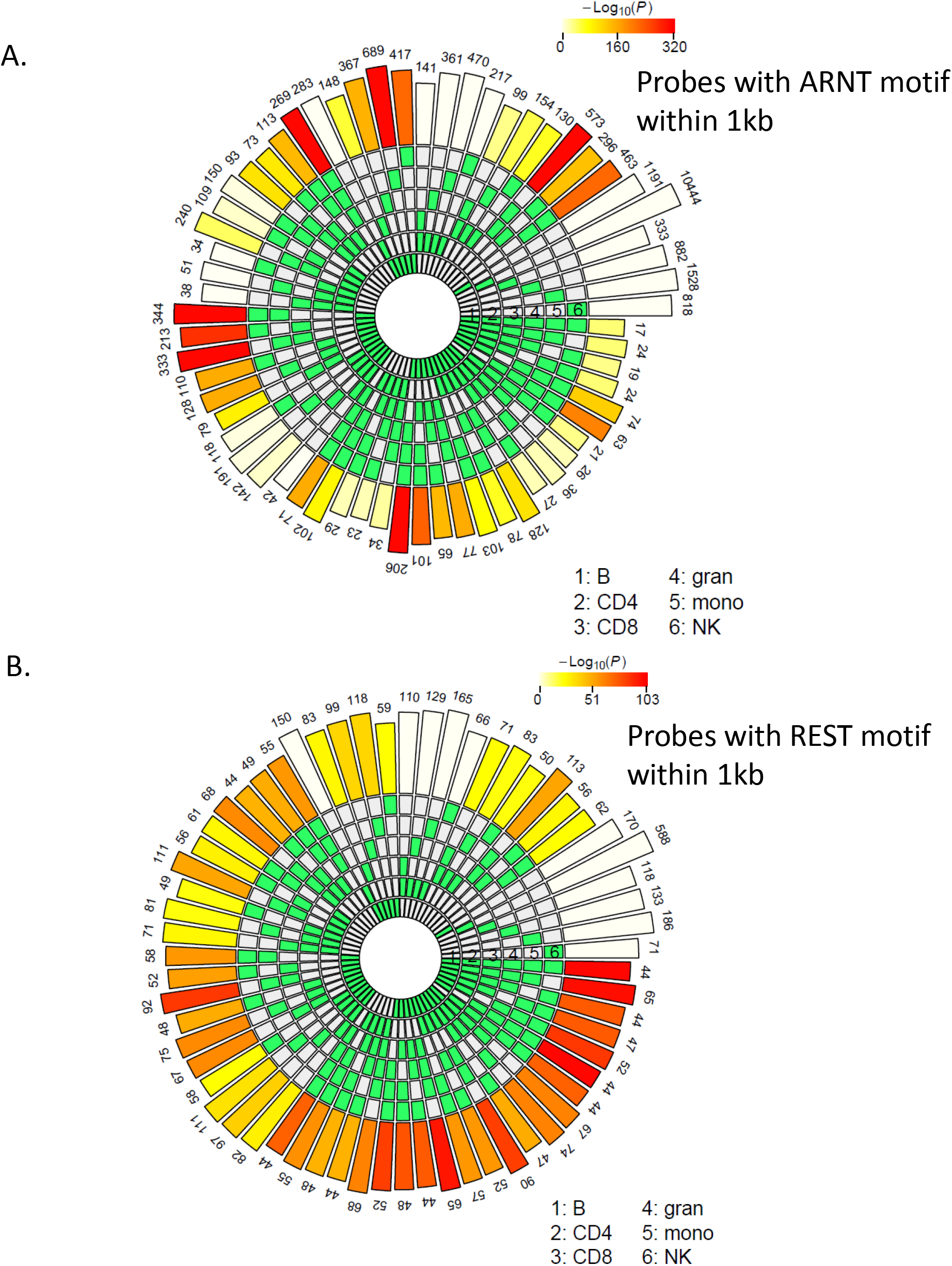
Count of age-associated hypo- or hypermethylated probes with ARNT or REST motifs within 1kb respectively. ARNT and REST were the top TF motifs associated respectively with all hypo- and hypermethylated age-associated probes in most cell types. To verify whether the same set of probes were present in each cell type with the ARNT(A) or REST(B) motifs, HOMER was used to find the probes that have a ARNT or REST motif within + 500bp. The SuperExact test based circular plots shows the overlap between the different cell types (relevant cell types are indicated in green boxes in the inner circles for each combination). The numbers above the outermost bars indicate the count of probes that have ARNT or REST motifs across various combinations of cells while the color of the outermost bars in the plot indicate the log transformed p values obtained from hypergeometric test to check whether the overlap is significant or not.

## LIST OF SUPPLMENTARY TABLES

**Supplementary Table 1:**
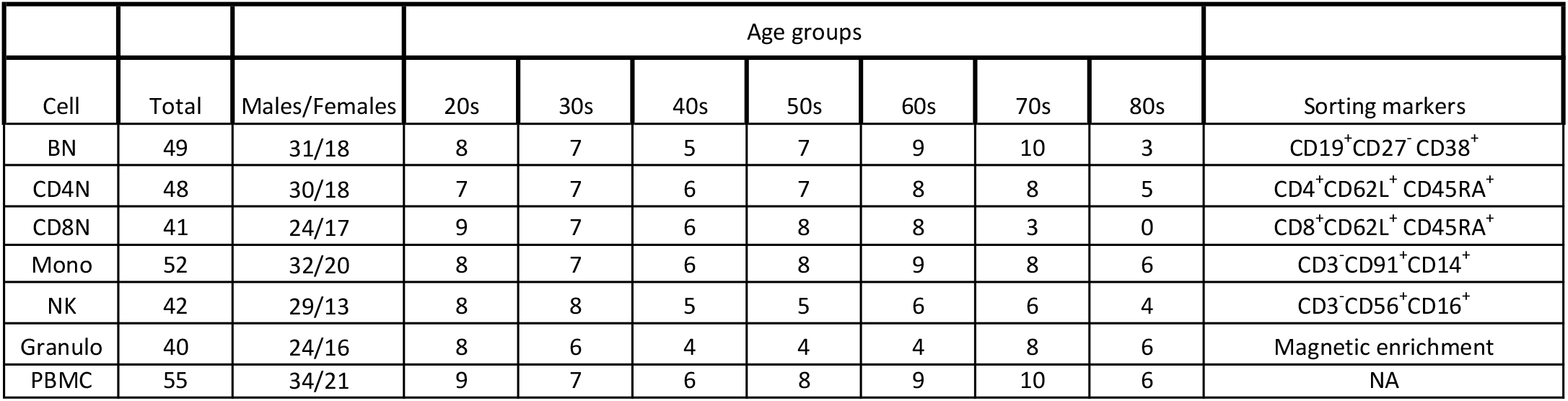
Demographic and flow cytometry marker details of the cohort. Details of the age and sex distribution of the healthy donors from the GESTALT study for each of the primary immune cell type population are described. The flow cytometry markers for cell selection are also mentioned.

**Supplementary Table 2:** Distribution of slope for probes significantly changing with age in the immune cells. The age-associated probes were identified from beta regression (FDR p<0.05).

|  | BN |  | CD4N |  | CD8N |  | granulocytes |  | monocytes |  | NK |  |
| --- | --- | --- | --- | --- | --- | --- | --- | --- | --- | --- | --- | --- |
| β-coefficient | hypoage* | hyperage* | hypoage | hyperage | hypoage | hyperage | hypoage | hyperage | hypoage | hyperage | hypoage | hyperage |
| <0.01 | 5418 | 3558 | 63185 | 40787 | 1446 | 1576 | 4077 | 1211 | 7344 | 5186 | 3509 | 506 |
| 0.01-0.02 | 1018 | 287 | 321 | 274 | 379 | 122 | 525 | 116 | 824 | 211 | 609 | 109 |
| 0.02-0.03 | 63 | 17 | 56 | 21 | 24 | 5 | 24 | 4 | 46 | 14 | 28 | 7 |
| >0.03 | 8 | 1 | 3 | 1 | 1 | 0 | 1 | 0 | 3 | 0 | 2 | 1 |
| Total | 6507 | 3863 | 63565 | 41083 | 1850 | 1703 | 4627 | 1331 | 8217 | 5411 | 4148 | 623 |

hypoage\*- probes hypomethylated with age
hyperage\*- probes hypermethylated with age

**Supplementary Table 3:**
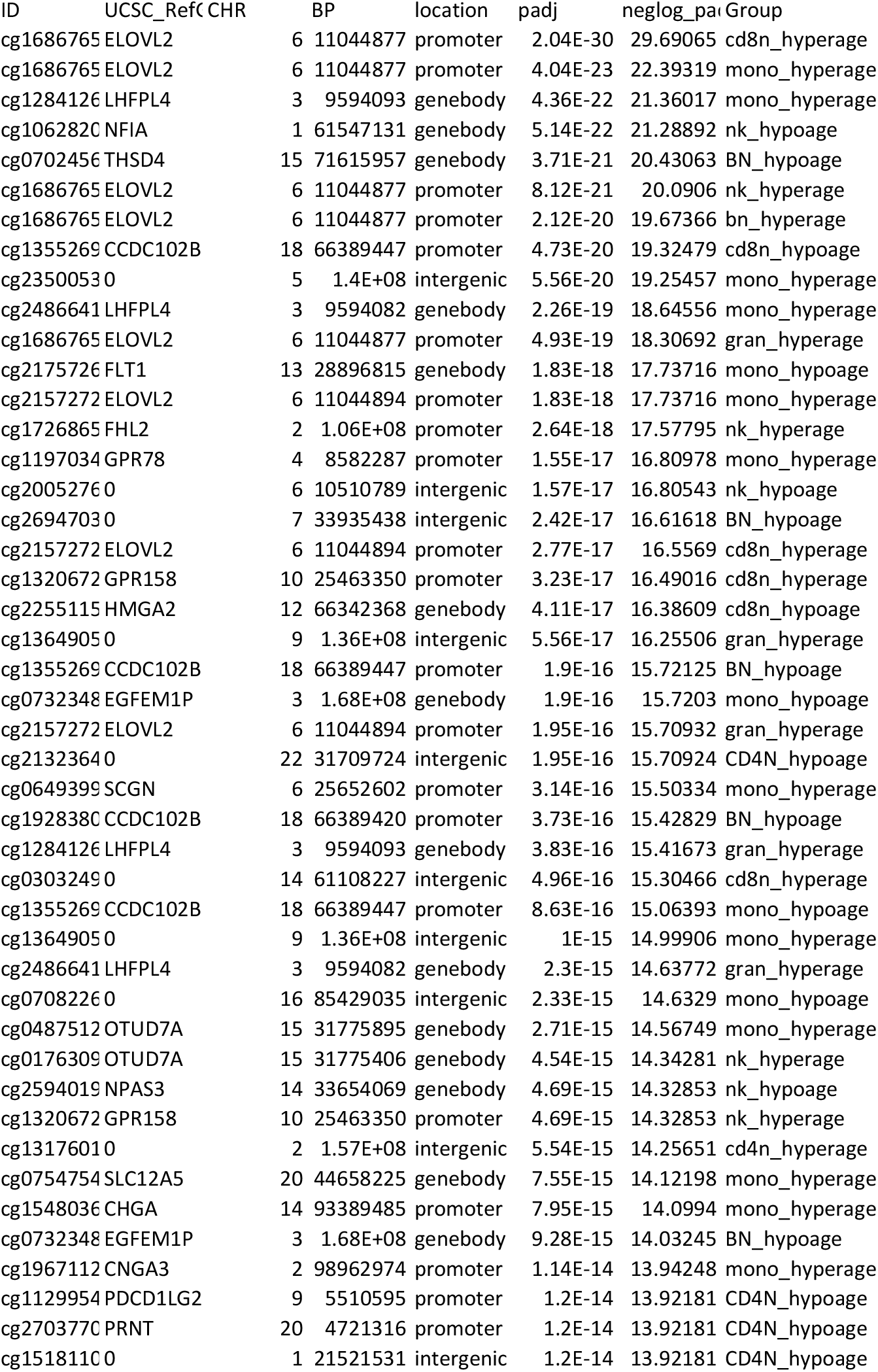
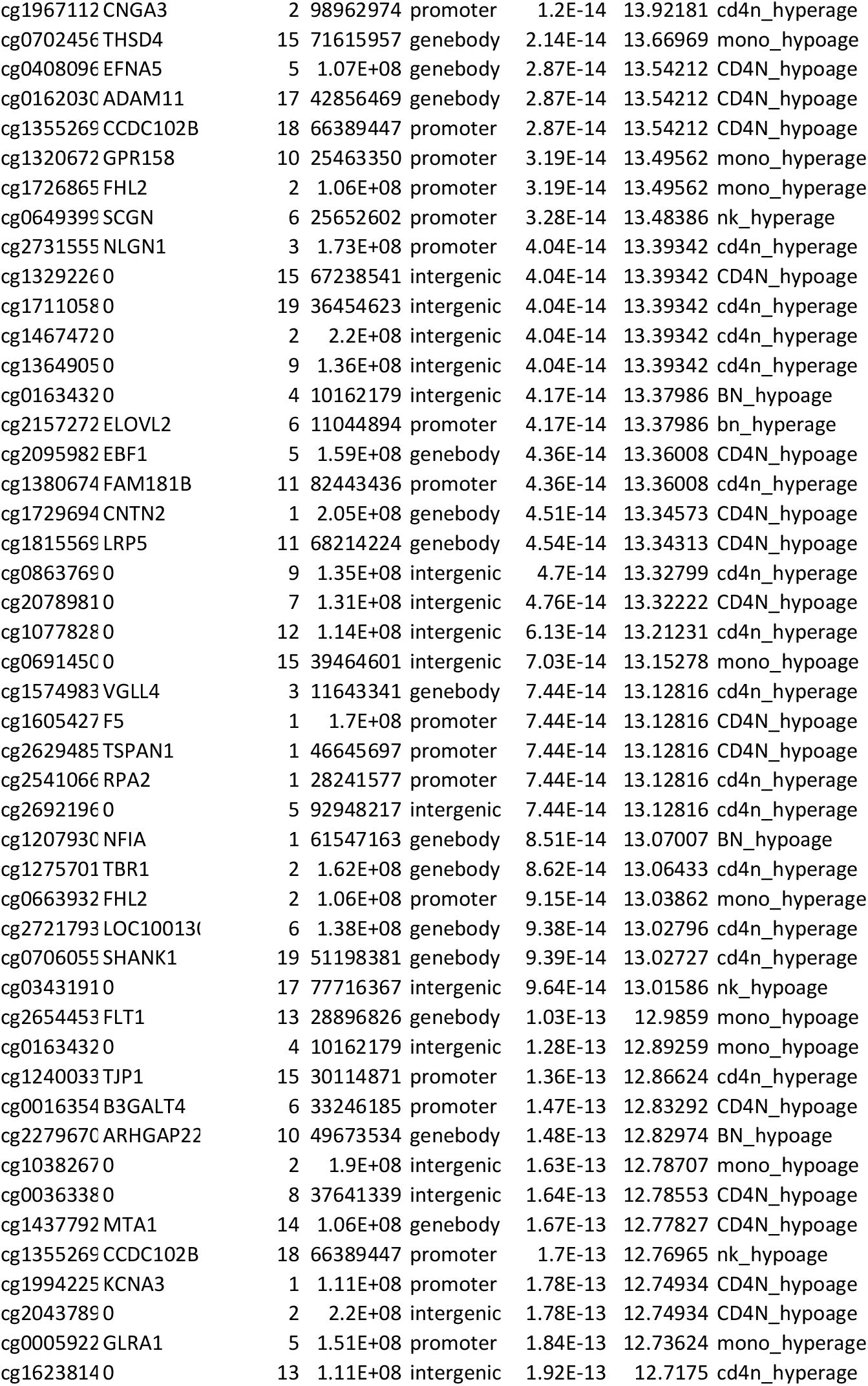
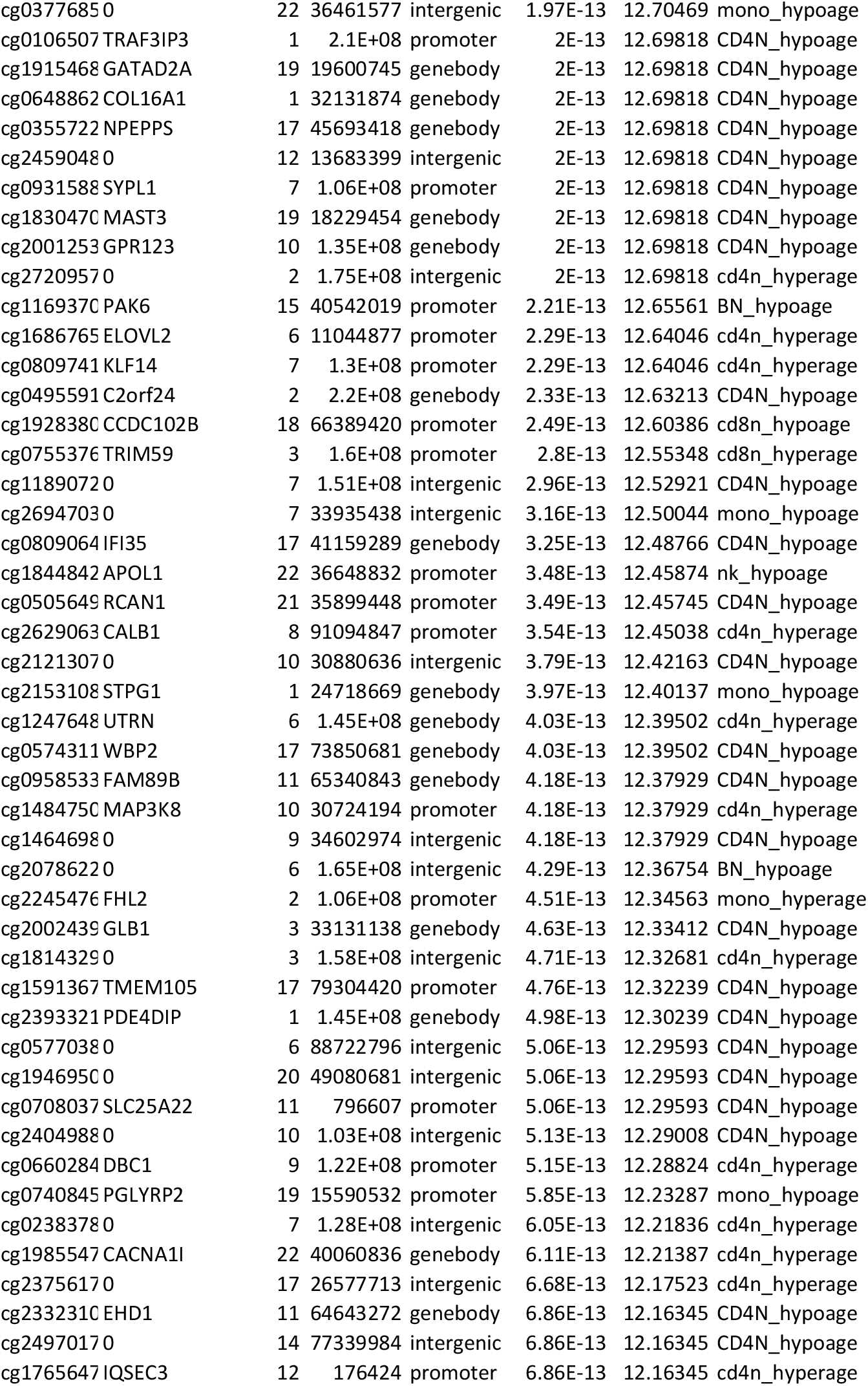
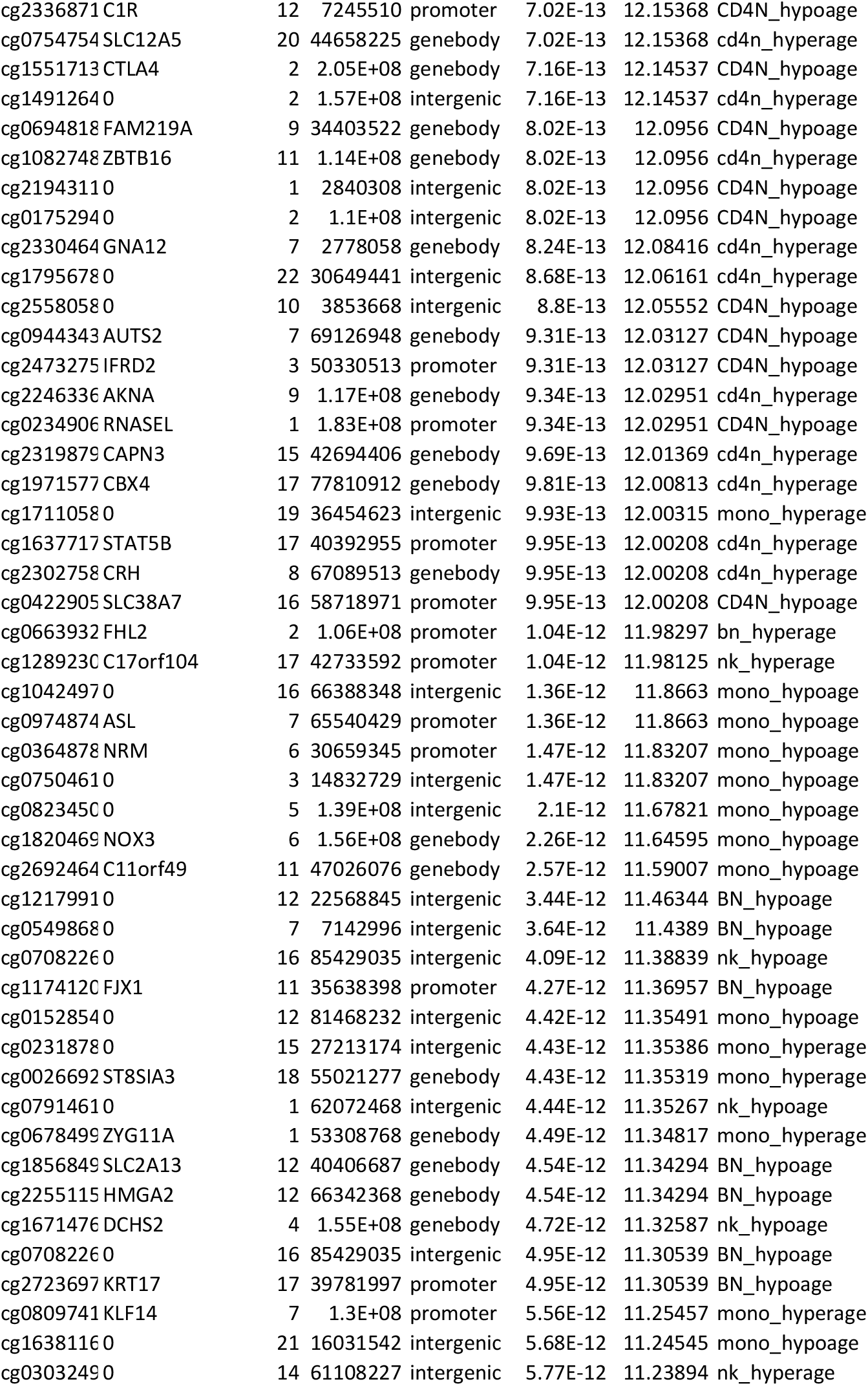
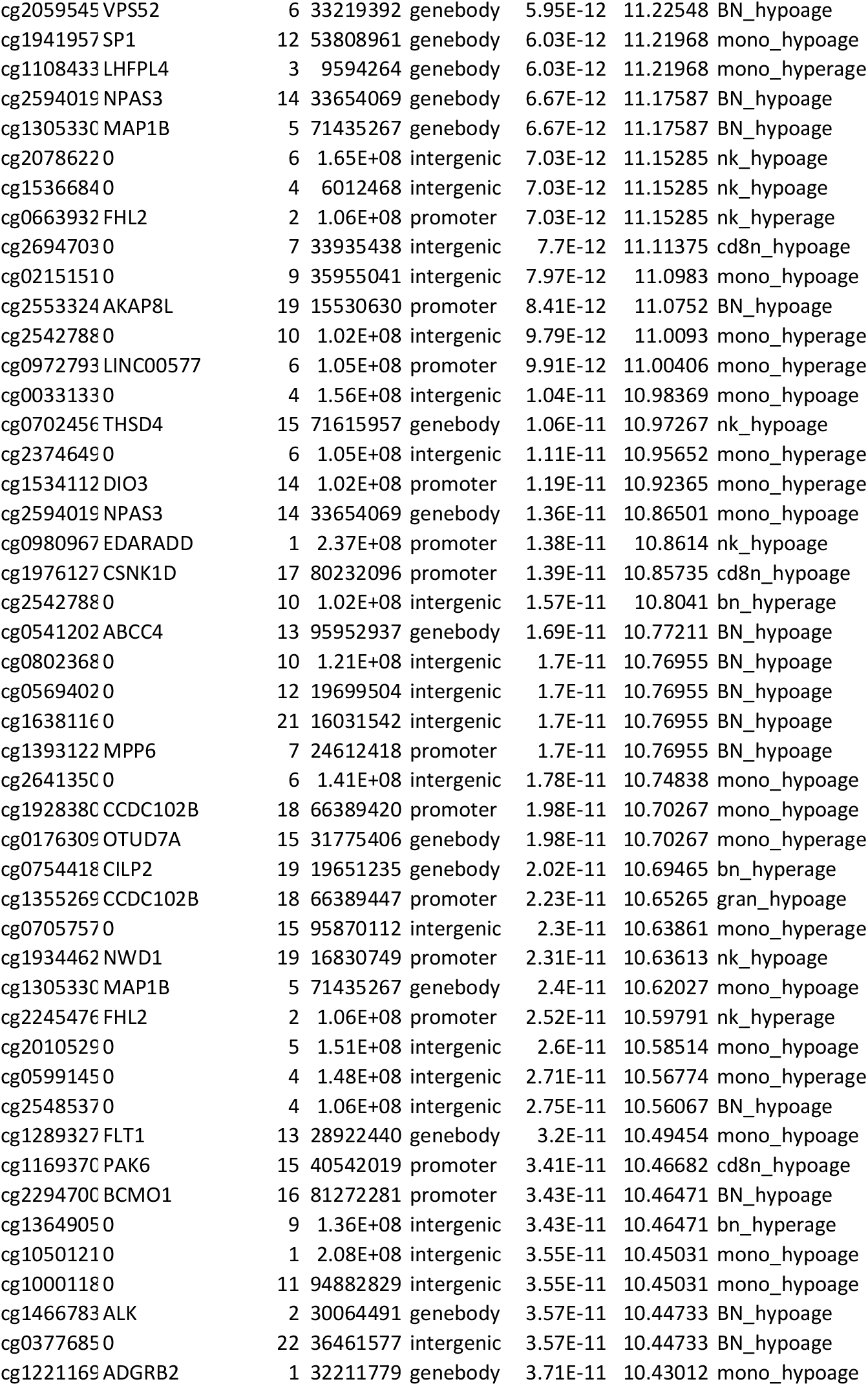
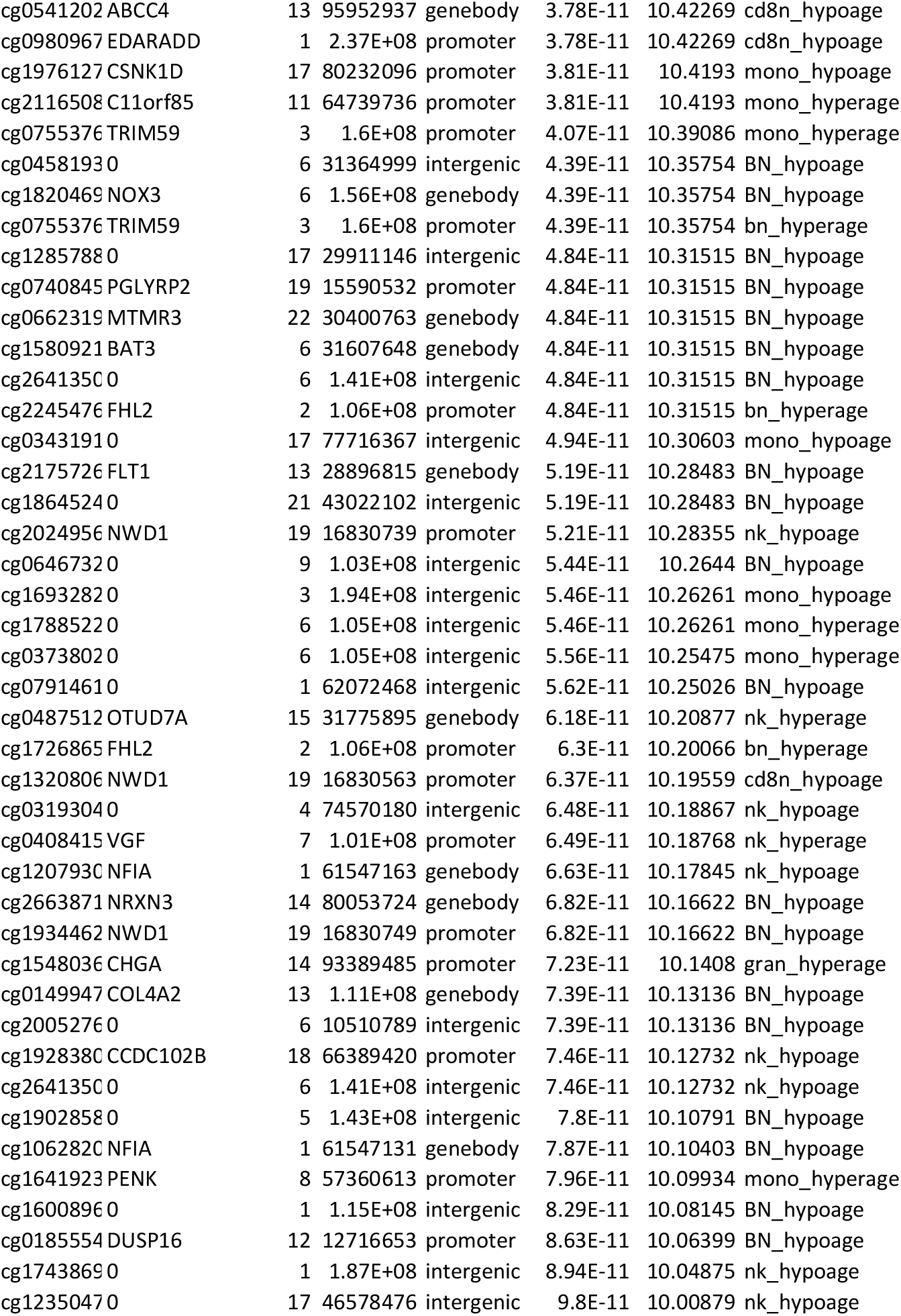
List of most significant age-associated probes in the immune cells. Based on a p-value cut off (-log (FDR adjusted p) >10), the top age-associated candidates were studied to search for common genes across all immune cells.

**Supplementary Table 4:**
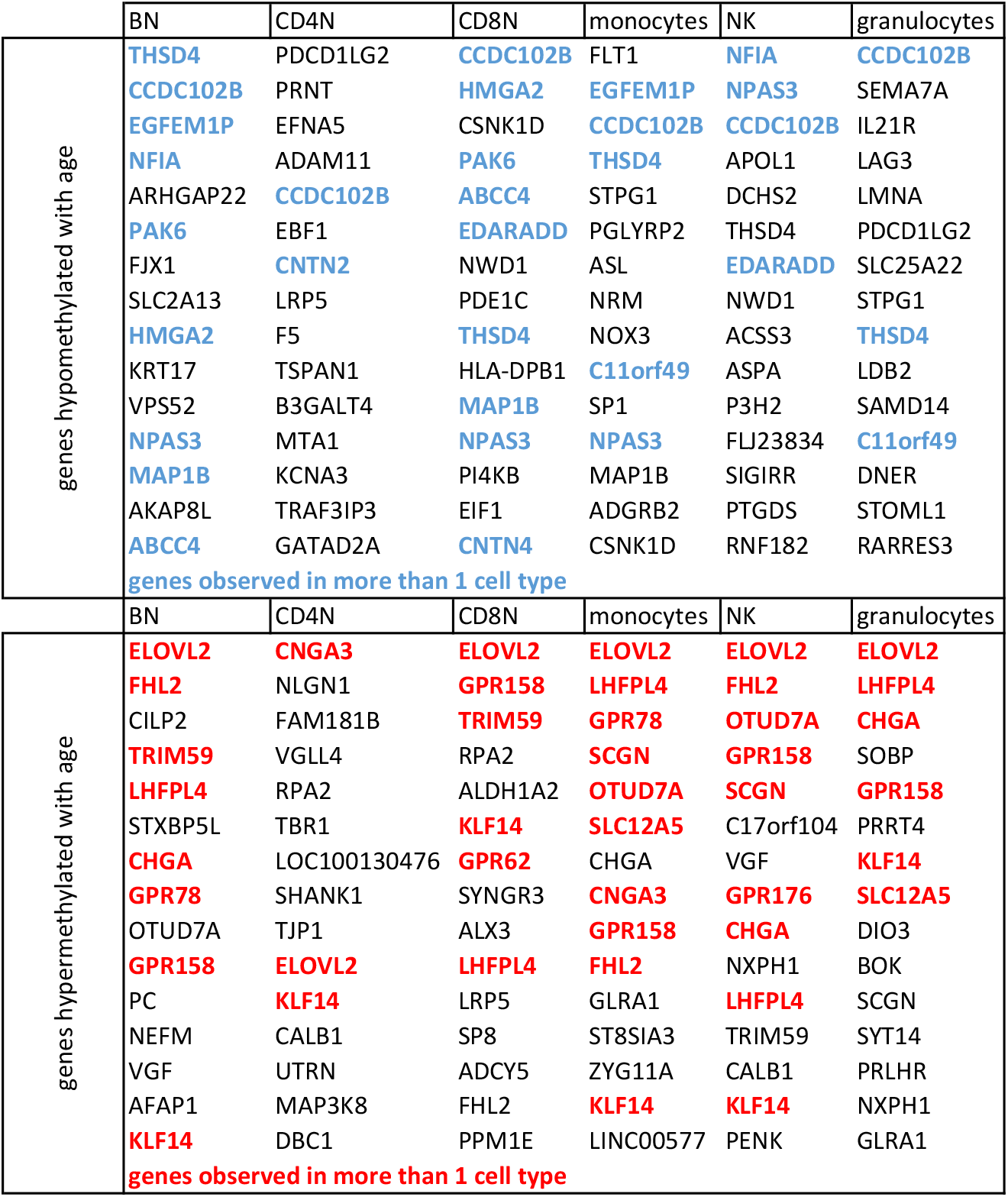
List of top age-associated genes in the six immune cell types. The list of genes from top 50 age-associated hypo- and hypermethylated probes.

**Supplementary Table 5:**
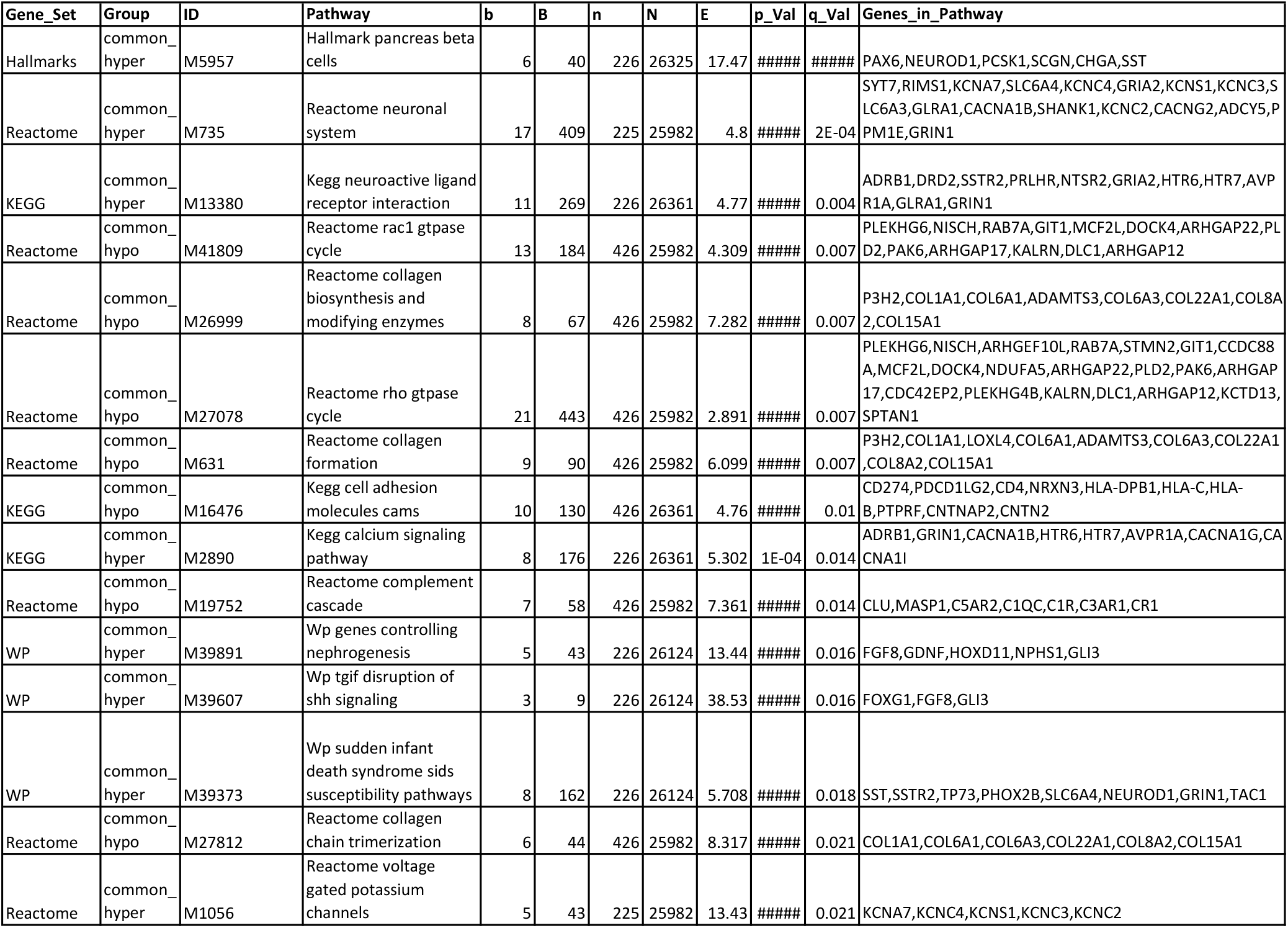
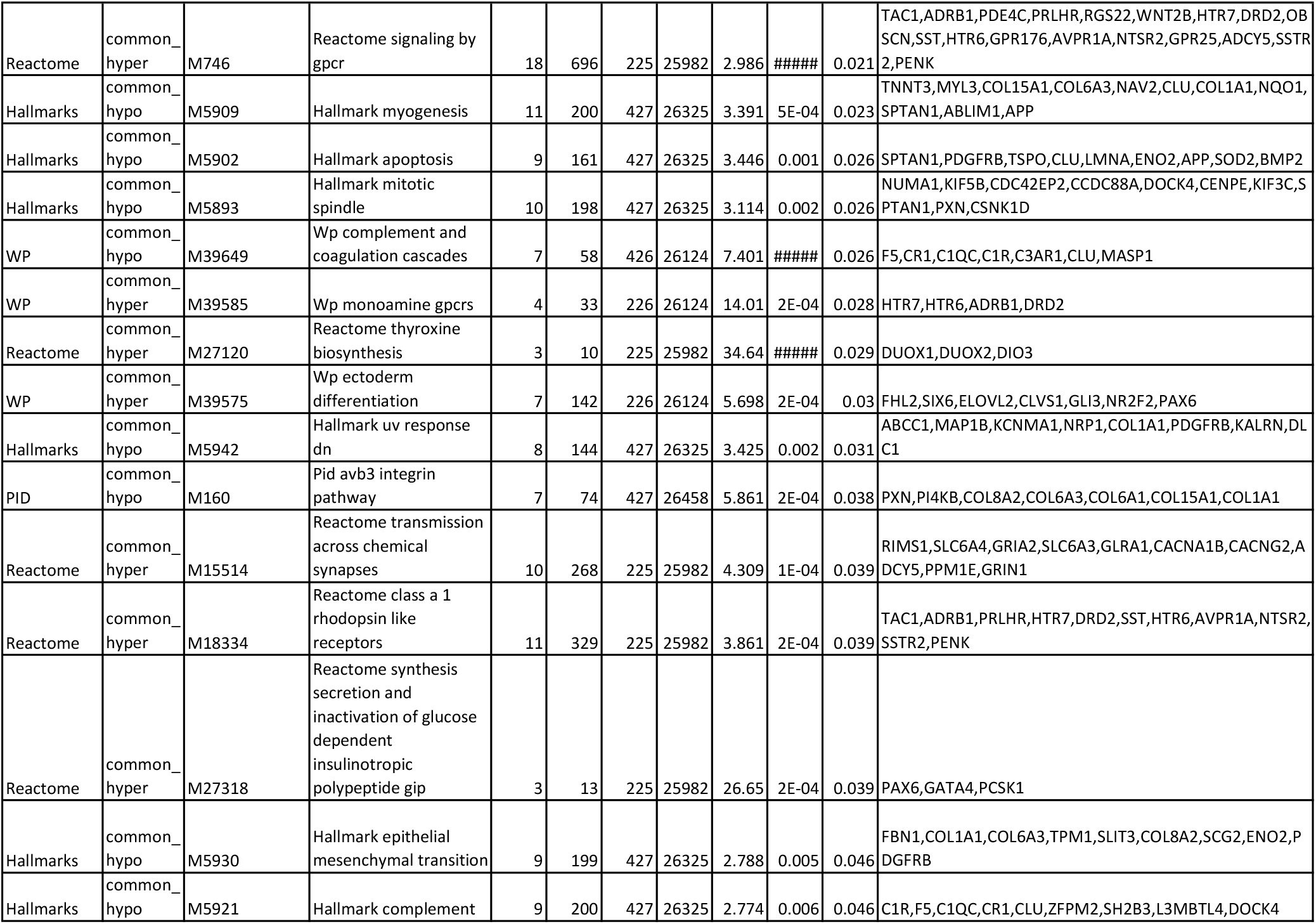
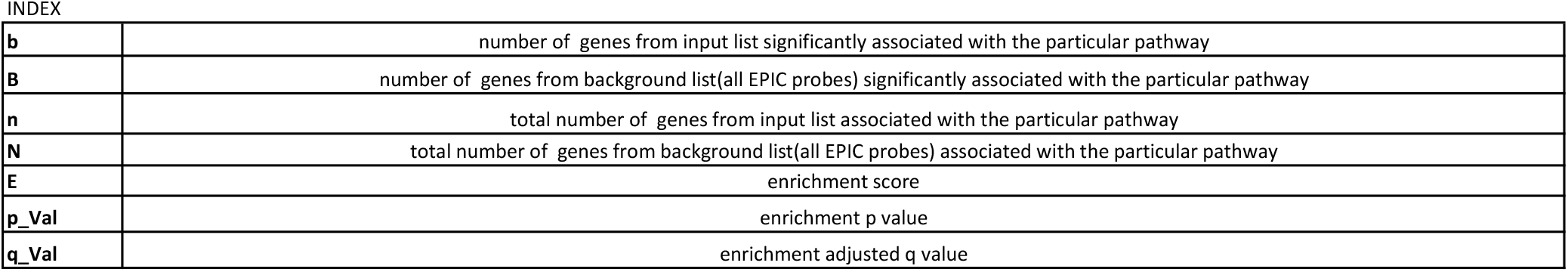
Detailed output of Gene Set Enrichment Analysis. Gene Set Enrichment Analysis was performed on genes based on annotation of age-associated hypo- and hypermethylation probes commonly changing in 5 or more cell types.

| Gene_Set | Group | ID | Pathway | b | B | n | N | E | p_Val | q_Val | Genes_in_Pathway |
| --- | --- | --- | --- | --- | --- | --- | --- | --- | --- | --- | --- |
| Hallmarks | common_<br>hyper | M5957 | Hallmark pancreas beta cells | 6 | 40 | 226 | 26325 | 17.47 | ##### | ##### | PAX6,NEUROD1,PCSK1,SCGN,CHGA,SST |
| Reactome | common_<br>hyper | M735 | Reactome neuronal system | 17 | 409 | 225 | 25982 | 4.8 | ##### | 2E-04 | SYT7,RIMS1,KCNA7,SLC6A4,KCNC4,GRIA2,KCNS1,KCNC3,SLC6A3,GLRA1,CACNA1B,SHANK1,KCNC2,CACNG2,ADCY5,PPM1E,GRIN1 |
| KEGG | common_<br>hyper | M13380 | Kegg neuroactive ligand receptor interaction | 11 | 269 | 226 | 26361 | 4.77 | ##### | 0.004 | ADRB1,DRD2,SSTR2,PRLHR,NTSR2,GRIA2,HTR6,HTR7,AVPR1A,GLRA1,GRIN1 |
| Reactome | common_<br>hypo | M41809 | Reactome rac1 gtpase cycle | 13 | 184 | 426 | 25982 | 4.309 | ##### | 0.007 | PLEKHG6,NISCH,RAB7A,GIT1,MCF2L,DOCK4,ARHGAP22,PLD2,PAK6,ARHGAP17,KALRN,DLC1,ARHGAP12 |
| Reactome | common_<br>hypo | M26999 | Reactome collagen biosynthesis and modifying enzymes | 8 | 67 | 426 | 25982 | 7.282 | ##### | 0.007 | P3H2,COL1A1,COL6A1,ADAMTS3,COL6A3,COL22A1,COL8A2,COL15A1 |
| Reactome | common_<br>hypo | M27078 | Reactome rho gtpase cycle | 21 | 443 | 426 | 25982 | 2.891 | ##### | 0.007 | PLEKHG6,NISCH,ARHGEF10L,RAB7A,STMN2,GIT1,CCDC88A,MCF2L,DOCK4,NDUFA5,ARHGAP22,PLD2,PAK6,ARHGAP17,CDC42EP2,PLEKHG4B,KALRN,DLC1,ARHGAP12,KCTD13,SPTAN1 |
| Reactome | common_<br>hypo | M631 | Reactome collagen formation | 9 | 90 | 426 | 25982 | 6.099 | ##### | 0.007 | P3H2,COL1A1,LOXL4,COL6A1,ADAMTS3,COL6A3,COL22A1,COL8A2,COL15A1 |
| KEGG | common_<br>hypo | M16476 | Kegg cell adhesion molecules cams | 10 | 130 | 426 | 26361 | 4.76 | ##### | 0.01 | CD274,PDCD1LG2,CD4,NRXN3,HLA-DPB1,HLA-C,HLA-B,PTPRF,CNTNAP2,CNTN2 |
| KEGG | common_<br>hyper | M2890 | Kegg calcium signaling pathway | 8 | 176 | 226 | 26361 | 5.302 | 1E-04 | 0.014 | ADRB1,GRIN1,CACNA1B,HTR6,HTR7,AVPR1A,CACNA1G,CACNA1I |
| Reactome | common_<br>hypo | M19752 | Reactome complement cascade | 7 | 58 | 426 | 25982 | 7.361 | ##### | 0.014 | CLU,MASP1,C5AR2,C1QC,C1R,C3AR1,CR1 |
| WP | common_<br>hyper | M39891 | Wp genes controlling nephrogenesis | 5 | 43 | 226 | 26124 | 13.44 | ##### | 0.016 | FGF8,GDNF,HOXD11,NPHS1,GLI3 |
| WP | common_<br>hyper | M39607 | Wp tgif disruption of shh signaling | 3 | 9 | 226 | 26124 | 38.53 | ##### | 0.016 | FOXG1,FGF8,GLI3 |
| WP | common_<br>hyper | M39373 | Wp sudden infant death syndrome sids susceptibility pathways | 8 | 162 | 226 | 26124 | 5.708 | ##### | 0.018 | SST,SSTR2,TP73,PHOX2B,SLC6A4,NEUROD1,GRIN1,TAC1 |
| Reactome | common_<br>hypo | M27812 | Reactome collagen chain trimerization | 6 | 44 | 426 | 25982 | 8.317 | ##### | 0.021 | COL1A1,COL6A1,COL6A3,COL22A1,COL8A2,COL15A1 |
| Reactome | common_<br>hyper | M1056 | Reactome voltage gated potassium channels | 5 | 43 | 225 | 25982 | 13.43 | ##### | 0.021 | KCNA7,KCNC4,KCNS1,KCNC3,KCNC2 |
| Reactome | common_hyper | M746 | Reactome signaling by gpcr | 18 | 696 | 225 | 25982 | 2.986 | ##### | 0.021 | TAC1,ADRB1,PDE4C,PRLHR,RGS22,WNT2B,HTR7,DRD2,OBSCN,SST,HTR6,GPR176,AVPR1A,NTSR2,GPR25,ADCY5,SSTR2,PENK |
| Hallmarks | common_hypo | M5909 | Hallmark myogenesis | 11 | 200 | 427 | 26325 | 3.391 | 5E-04 | 0.023 | TNNT3,MYL3,COL15A1,COL6A3,NAV2,CLU,COL1A1,NQO1,SPTAN1,ABLIM1,APP |
| Hallmarks | common_hypo | M5902 | Hallmark apoptosis | 9 | 161 | 427 | 26325 | 3.446 | 0.001 | 0.026 | SPTAN1,PDGFRB,TSPO,CLU,LMNA,ENO2,APP,SOD2,BMP2 |
| Hallmarks | common_hypo | M5893 | Hallmark mitotic spindle | 10 | 198 | 427 | 26325 | 3.114 | 0.002 | 0.026 | NUMA1,KIF5B,CDC42EP2,CCDC88A,DOCK4,CENPE,KIF3C,SPTAN1,PXN,CSNK1D |
| WP | common_hypo | M39649 | Wp complement and coagulation cascades | 7 | 58 | 426 | 26124 | 7.401 | ##### | 0.026 | F5,CR1,C1QC,C1R,C3AR1,CLU,MASP1 |
| WP | common_hyper | M39585 | Wp monoamine gpcrs | 4 | 33 | 226 | 26124 | 14.01 | 2E-04 | 0.028 | HTR7,HTR6,ADRB1,DRD2 |
| Reactome | common_hyper | M27120 | Reactome thyroxine biosynthesis | 3 | 10 | 225 | 25982 | 34.64 | ##### | 0.029 | DUOX1,DUOX2,DIO3 |
| WP | common_hyper | M39575 | Wp ectoderm differentiation | 7 | 142 | 226 | 26124 | 5.698 | 2E-04 | 0.03 | FHL2,SIX6,ELOVL2,CLVS1,GLI3,NR2F2,PAX6 |
| Hallmarks | common_hypo | M5942 | Hallmark uv response dn | 8 | 144 | 427 | 26325 | 3.425 | 0.002 | 0.031 | ABCC1,MAP1B,KCNMA1,NRP1,COL1A1,PDGFRB,KALRN,DL C1 |
| PID | common_hypo | M160 | Pid avb3 integrin pathway | 7 | 74 | 427 | 26458 | 5.861 | 2E-04 | 0.038 | PXN,PI4KB,COL8A2,COL6A3,COL6A1,COL15A1,COL1A1 |
| Reactome | common_hyper | M15514 | Reactome transmission across chemical synapses | 10 | 268 | 225 | 25982 | 4.309 | 1E-04 | 0.039 | RIMS1,SLC6A4,GRIA2,SLC6A3,GLRA1,CACNA1B,CACNG2,ADCY5,PPM1E,GRIN1 |
| Reactome | common_hyper | M18334 | Reactome class a 1 rhodopsin like receptors | 11 | 329 | 225 | 25982 | 3.861 | 2E-04 | 0.039 | TAC1,ADRB1,PRLHR,HTR7,DRD2,SST,HTR6,AVPR1A,NTSR2,SSTR2,PENK |
| Reactome | common_hyper | M27318 | Reactome synthesis secretion and inactivation of glucose dependent insulinotropic polypeptide gip | 3 | 13 | 225 | 25982 | 26.65 | 2E-04 | 0.039 | PAX6,GATA4,PCSK1 |
| Hallmarks | common_hypo | M5930 | Hallmark epithelial mesenchymal transition | 9 | 199 | 427 | 26325 | 2.788 | 0.005 | 0.046 | FBN1,COL1A1,COL6A3,TPM1,SLIT3,COL8A2,SCG2,ENO2,PDGFRB |
| Hallmarks | common_hypo | M5921 | Hallmark complement | 9 | 200 | 427 | 26325 | 2.774 | 0.006 | 0.046 | C1R,F5,C1QC,CR1,CLU,ZFPM2,SH2B3,L3MBTL4,DOCK4 |

INDEX
|  |  |
| --- | --- |
| **b** | number of genes from input list significantly associated with the particular pathway |
| **B** | number of genes from background list(all EPIC probes) significantly associated with the particular pathway |
| **n** | total number of genes from input list associated with the particular pathway |
| **N** | total number of genes from background list(all EPIC probes) associated with the particular pathway |
| **E** | enrichment score |
| **p_Val** | enrichment p value |
| **q_Val** | enrichment adjusted q value |

**Supplementary Table 6:**
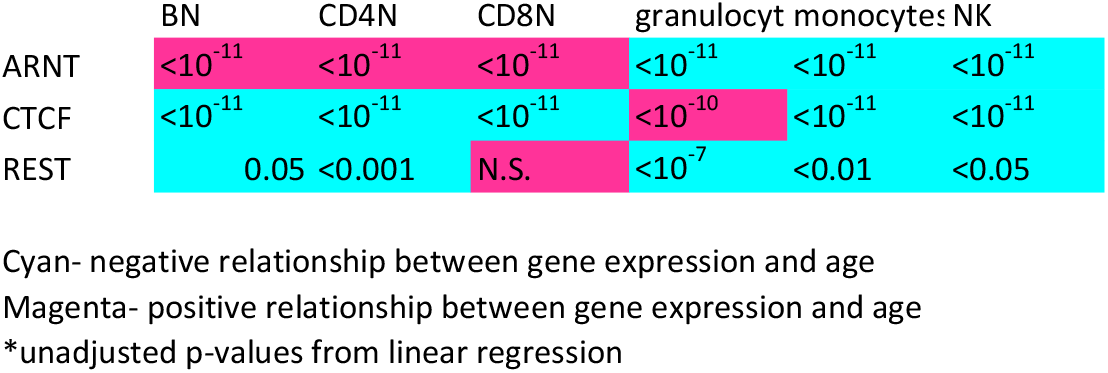
Age-associated differences of transcripts for ARNT, REST and CTCF. RNASeq data was used to look into the gene expression change of the selected transcription factors with age. These transcription factor motifs are most commonly associated with the age-related methylated sites in all immune cells.

